# The endoplasmic reticulum adopts two distinct tubule forms

**DOI:** 10.1101/2021.10.26.465969

**Authors:** Bowen Wang, Zhiheng Zhao, Michael Xiong, Rui Yan, Ke Xu

## Abstract

The endoplasmic reticulum (ER) is a versatile organelle with diverse functions. Through super-resolution microscopy, we show that the peripheral ER in the mammalian cell adopts two distinct forms of tubules. Whereas an ultrathin form, R1, is consistently covered by ER-membrane curvature-promoting proteins, *e*.*g*., Rtn4 in the native cell, in the second form, R2, Rtn4 and analogs are arranged into two parallel lines at a conserved separation of ∼105 nm over long ranges. The two tubule forms together account for ∼90% of the total tubule length in the cell, with either one being dominant in different cell types. The R1-R2 dichotomy and the final tubule geometry are both co-regulated by Rtn4 (and analogs) and the ER sheet-maintaining protein Climp63, which respectively define the edge curvature and lumen height of the R2 tubules to generate a ribbon-like structure of well-defined width. Accordingly, the R2 tubule width correlates positively with the Climp63 intralumenal size. The R1 and R2 tubules undergo active remodeling at the second/sub-second time scales as they differently accommodate proteins, with the former effectively excluding ER-luminal proteins and ER-membrane proteins with large intraluminal domains. We thus uncover a dynamic structural dichotomy for ER tubules with intriguing functional implications.

## Introduction

Being the largest and most expansive organelle in the cell, the endoplasmic reticulum (ER) carries diverse key functions from protein and lipid synthesis, protein folding and modification, transport, calcium storage, to organelle interactions^1-6^. The shaping mechanism of this complex, membrane-bounded organelle is thus of fundamental significance^7-21^.

The fully connected ER system is classically subdivided into three distinct domains, namely the nuclear envelope, ER sheets, and ER tubules. The latter two structures, collectively known as the peripheral ER, interweave into a dynamic, interconvertible network, yet are differently maintained and regulated by ER-shaping proteins. In mammalian cells, the two-dimensional ER sheets consist of two flat lipid bilayers at an ∼50 nm separation^3,4,11^; this conserved luminal spacing is maintained by Climp63 (CKAP4), a transmembrane protein that forms intraluminal bridges between the opposing membranes via its extended coiled-coil domain^11,20^. Meanwhile, the highly curved edges of the ER sheets are stabilized by curvature-promoting proteins including the reticulons (Rtns) and REEPs^8,11^, which, with their hairpin-like topology, insert into the outer leaflet of the lipid bilayer as wedges^9,10,12,21^. The same curvature-promoting proteins also stabilize the high curvatures in the one-dimensional ER tubules^9,10^, which in mammalian cells are often taken as cylindrical tubules of ∼50-100 nm diameter^2,7^. Notably, the intracellular overexpression of the reticulon Rtn4 (Nogo) substantially reduces the ER tubule diameter to ∼20 nm, highlighting its ability to shape high-curvature tubules^10^.

The rise of super-resolution microscopy (SRM) over the past decade offers new means to discover cellular structures^22-24^. When applied to the ER^14,17-19,25,26^, new perspectives have emerged to challenge the traditional division between ER sheets and tubules, *e*.*g*., whether the peripheral ER sheets should be viewed as a matrix of tubules or sheets with many nanoscale holes^17,19^. Here, we instead focus on the ER tubules, and, unexpectedly, suggest that a substantial fraction of the ER tubules should be recognized as thinned and elongated sheets of fixed widths defined by Climp63-Rtn interactions.

## Results

### STORM unveils an ER-tubule dichotomy

We started by applying stochastic optical reconstruction microscopy (STORM) SRM^27,28^, which routinely achieves ∼20 nm spatial resolution, to immunolabeled endogenous Rtn4 in untransfected COS-7 cells. Rtn4 is one of the most studied ER-membrane curvature-promoting proteins in the mammalian cell, and quantitative proteomics of a human cell line has indicated it as the most abundant member of the group^29^. Similar results were obtained using two different antibodies (Fig. S1), and immunoblotting indicated that Rtn4b and Rtn4b2 were the dominant Rtn4 forms detected in our experiments (Fig. S2).

Unexpectedly, we found that in a major fraction of the ER tubules, Rtn4 showed up as two parallel lines over long ranges (filled arrowheads in Fig. 1ab). The typical center-to-center distance between the two lines was ∼105 nm (Fig. 1c), and statistics showed a conserved narrow distribution of 106±18 nm between different cell types (Fig. 1d). This well-defined separation suggests a distinct structural arrangement and hence classification, which we hereby designate as R2 (Rtn double-line).

**Fig. 1.**
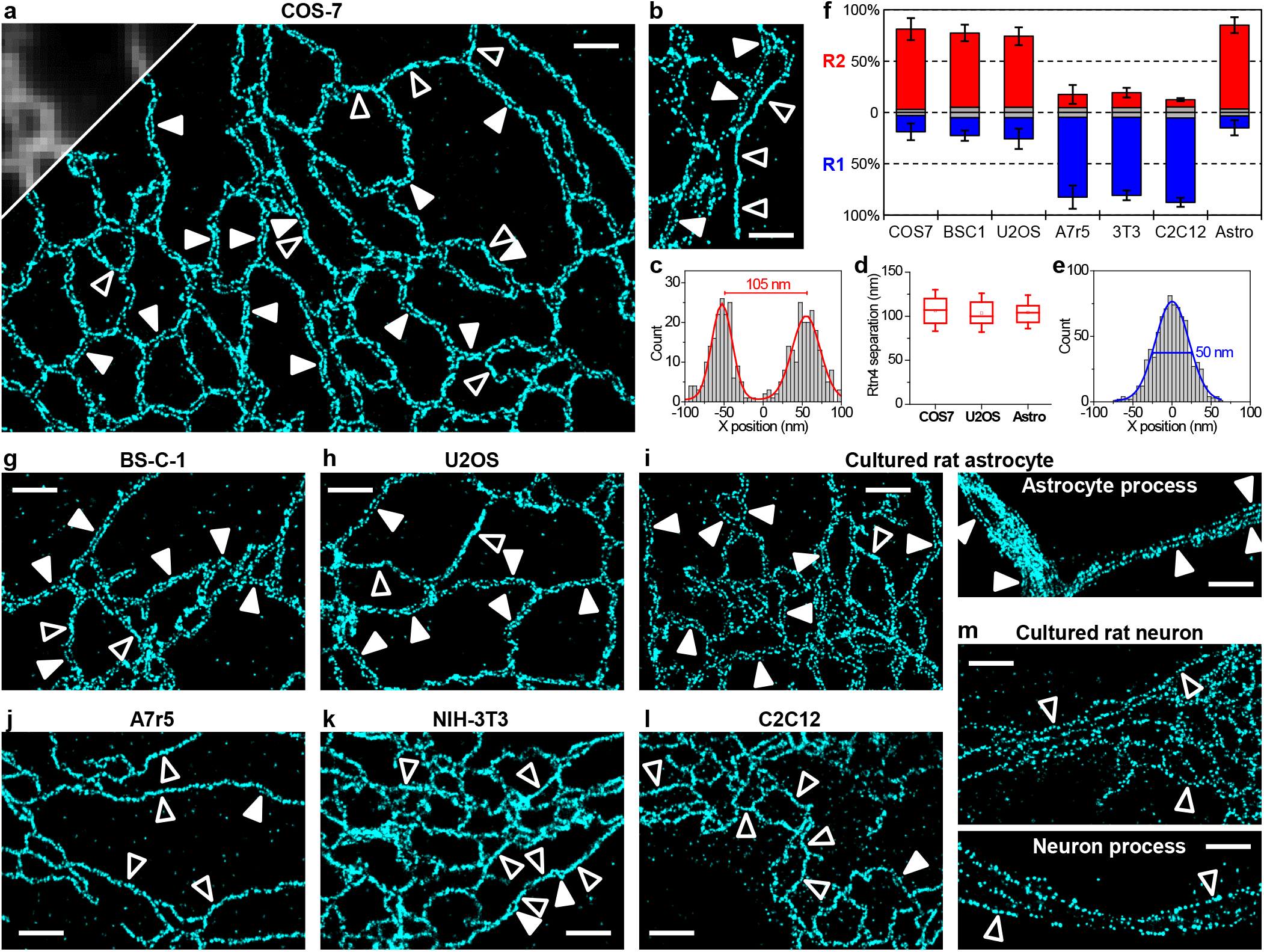
STORM SRM of endogenous Rtn4 unveils a structural dichotomy for ER tubules. (**a**) Representative STORM image of immunolabeled Rtn4 in the untransfected COS-7 cell, compared to diffraction-limited fluorescence image (upper-left corner). Filled and hollow arrowheads point to examples of R2 (Rtn double-line) and R1 (Rtn single-line) tubules, respectively. (**b**) Result from another COS-7 cell, highlighting a long R1 tubule at the extremity of the ER network. (**c**) Representative STORM image intensity (as single-molecule counts) across an R2 tubule. (**d**) Distribution of the STORM-determined center-to-center separations between opposing Rtn4 lines in the R2 tubules in the native COS-7 cells, U2OS cells, and cultured rat astrocytes. Whiskers and boxes show 10%, 25%, 50%, 75%, and 90% quantiles. For each data point, *n* = 200 local distances pooled from 5 cells. (**e**) Representative STORM image intensity across an R1 tubule. Blue curve: fit to a gaussian distribution of 50 nm FWHM. (**f**) The fractions of R1 and R2 tubules by length, observed in different cell types. Blue: R1; red: R2; gray: ambiguous. Error bars: standard deviations between *n* = 5 cells of each type. (**g**-**m**) Representative STORM images of the immunolabeled endogenous Rtn4 in different cell lines: BS-C-1 (monkey epithelial; g), U2OS (human epithelial; h), A7r5 (rat fibroblast; j), NIH-3T3 (mouse fibroblast; k), and C2C12 (mouse myoblast; l), as well as in cultured primary astrocytes (i) and neurons (m) from the rat hippocampus. Scale bars: 1 µm.

Meanwhile, we also observed a second class of ER tubules in which Rtn4 appeared as a thin single line (henceforth R1) of ∼50 nm FWHM (full width at half maximum) (Fig. 1e). In COS-7 cells, they existed as short segments connecting the R2 tubules (hollow arrowheads in Fig. 1a), but also as longer tubules, often at the cell periphery, as the extremities of the ER network (hollow arrowheads in Fig. 1b). Together, ∼75% and ∼15% of the total ER tubule lengths in COS-7 cells were classified as the R2 and R1 forms, respectively, with the remaining ∼10% being ambiguous (Fig. 1f). Thus, STORM unveiled an R1-R2 dichotomy of ER tubules, with R2 being the dominating form in COS-7 cells.

As we next examined seven other cell types, we found two epithelial cell lines, BS-C-1 (Fig. 1g) and U2OS (Fig. 1h), similarly exhibited an R2-dominant R1-R2 dichotomy (Fig. 1f). In contrast, two fibroblast cell lines (A7r5 and NIH-3T3; Fig. 1jk) and a myoblast cell line (C2C12; Fig. 1l) had >75% of the ER tubules in the R1 form and ∼10% in the R2 form as sporadic, short segments (filled arrowheads in Fig. 1j-l). A drastic contrast was further noted between the cultured primary astrocytes and neurons from the rat hippocampus: whereas in the astrocytes the ER tubules were predominantly (∼80%) R2, and this form was maintained along the thin, elongated processes (Fig. 1i), most ER tubules in the neurons appeared R1 (Fig. 1m).

### Regulation of the R1-R2 dichotomy

Our unexpected observation of two distinct ER tubule forms begs structural explanations. For the R1 tubules, the STORM-measured ∼50 nm apparent FWHM is comparable to that found (51 nm) for the ∼25 nm-diameter microtubules under similar experimental conditions^28^ given the antibody sizes and localization precisions. To simplify the discussion and consider the ∼4 nm lipid-bilayer thickness, we take the outer, inner (luminal), and mean diameters of the R1 tubule as 24, 16, and 20 nm, respectively. These very small diameters and hence high membrane curvatures may be maintained by a consistent coverage of the curvature-promoting Rtn4 along the circumference. Indeed, as we overexpressed Rtn4b-GFP in COS-7 cells, STORM showed that accompanying the reduction of ER sheets and enhanced presence of ER tubules^9^, most ER tubules turned into the ultrathin R1 form (Fig. 2bc and Fig. S3), in stark contrast to the ∼75% R2 fraction in the untransfected cells (Fig. 1af). Nonetheless, sporadic R2 segments were occasionally spotted along the predominantly R1 tubules (filled arrowhead in Fig. 2c), and some cells retained more R2 tubules (Fig. S3), suggesting that the R1-R2 dichotomy is still the preferred configuration under Rtn4-overexpression.

**Fig. 2.**
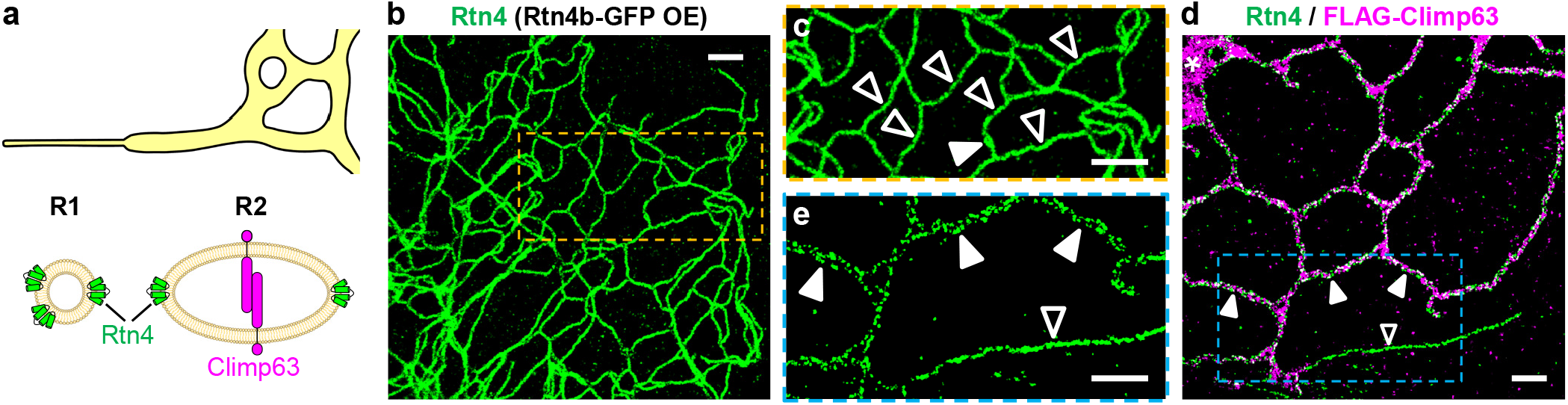
Rtn4 and Climp63 co-regulate the R1-R2 dichotomy. (**a**) Model: In the R1 tubules, high membrane curvatures are stabilized by a consistent Rtn4 coverage. In the R2 tubules, Rtn4 stabilizes the two highly curved edges of ribbon-like ER sheets, whereas Climp63 defines the luminal height. (**b**) Representative STORM image of immunolabeled Rtn4 in a COS-7 cell overexpressing Rtn4b-GFP. (**c**) Closeup of the box in (b). (**d**) Representative two-color STORM image of the immunolabeled endogenous Rtn4 (green) and overexpressed FLAG-Climp63 (magenta) in a COS-7 cell. Asterisk: ER sheet. (**e**) Closeup of the Rtn4 channel of the box in (d). Filled and hollow arrowheads point to examples of R2 and R1 tubules, respectively. Scale bars: 1 µm.

For the R2 tubules, the restriction of Rtn4 to two parallel lines raises a possibility that here Rtn4 stabilizes the edges of elongated, ribbon-like ER sheets (or compressed tubes) with two highly curved sides but relatively flat tops and bottoms (Fig. 2a). If that were the case, ER-sheet-maintaining proteins as Climp63 would likely regulate the luminal height of the R2 tubules as in typical ER sheets^11,20^. Indeed, fluorescence microscopy has shown the presence of Climp63 in ER tubules^11,17,18,30^, and a recent SRM study discusses the possible roles of Climp63 in regulating luminal compartmentalization and heterogeneity as “nanodomains”^18^. Two-color STORM showed that whereas Climp63 and Rtn4 respectively filled the ER sheets and delineated the sheet edges (asterisks in Fig. 2d and Fig. S4), as expected, a relatable structural arrangement further extended into the R2 tubules (filled arrowheads in Fig. 2de). In contrast, the R1 tubules were devoid of Climp63 (hollow arrowheads in Fig. 2de). High-level overexpression of Climp63 led to the expansion of ER sheets (Fig. S4)^11^, as well as an increased presence of R2 tubules in the otherwise R1-dominating NIH-3T3 cells (Fig. S4). Conversely, Climp63 siRNA markedly reduced R2 tubules in COS-7 cells (Fig. S5). Together, these results suggest Climp63 and Rtn4 co-regulate the R1-R2 dichotomy.

### Defining the R2 tubule width

To rationalize the remarkably conserved ∼105 nm distance between the opposing Rtn4 lines in the R2 tubules, we consider a simple model in which Climp63 sets the R2 tubule height *h* to the typical lumen height of ER sheets (∼50 nm)^11,20^, whereas Rtn4 holds the membrane radii of curvature at the two edges at some preferred value of *r*e. If the tubule adopts a smooth, elliptical cross-section, *r*e may be inferred from the simple ellipse geometry as *r*e = *h*^2^/2*w*, where *w* ∼100 nm is the ellipse width (∼105 nm Rtn4-Rtn4 separation minus lipid-bilayer thickness). For *h* ∼50 nm, *r*e is thus estimated as ∼12.5 nm, slightly larger than the assumed ∼10 nm radius of curvature of the ultrathin R1 tubules above, yet slightly smaller than the ∼15 nm value used in previous models for ER-sheet edges^11^.

Under the above simple model of fixed curvatures at the ribbon edges, the R2 tubule width *w* ∼*h*^2^/2*r*e would correlate positively with the lumen height *h* (Fig. 3a inset). Recent experiments have shown that for ER sheets, *h* can be altered by varying the length of the intraluminal coiled-coil domain of Climp63^20^. We utilized this strategy and STORM-imaged Rtn4 in COS-7 cells expressing different mCherry-tagged Climp63 variants. Expression of the full-length mouse Climp63(1-575) reaffirmed the respective presence and absence of Climp63 in the R2 and R1 tubules (Fig. 3b), with the STORM-determined Rtn4-Rtn4 separation in the former unchanged (107±17 nm; Fig. 3a). In contrast, the expression of a mutant that doubled the intraluminal domain, Climp63(2xlumen), which may raise *h* to ∼70 nm (ref ^20^), substantially increased the Rtn4-Rtn4 separation in R2 tubules to ∼155 nm (Fig. 3ac). Conversely, the expression of mutants with shortened intraluminal domains, Climp63(1-437), Climp63(1-336), and Climp63(1-301), led to progressive reductions of the Rtn4-Rtn4 separation (Fig. 3a,d-f), down to ∼75 nm for the shortest version (Fig. 3af). Together, our results indicate that Climp63 and Rtn4 jointly define the R2 tubule width.

**Fig. 3.**
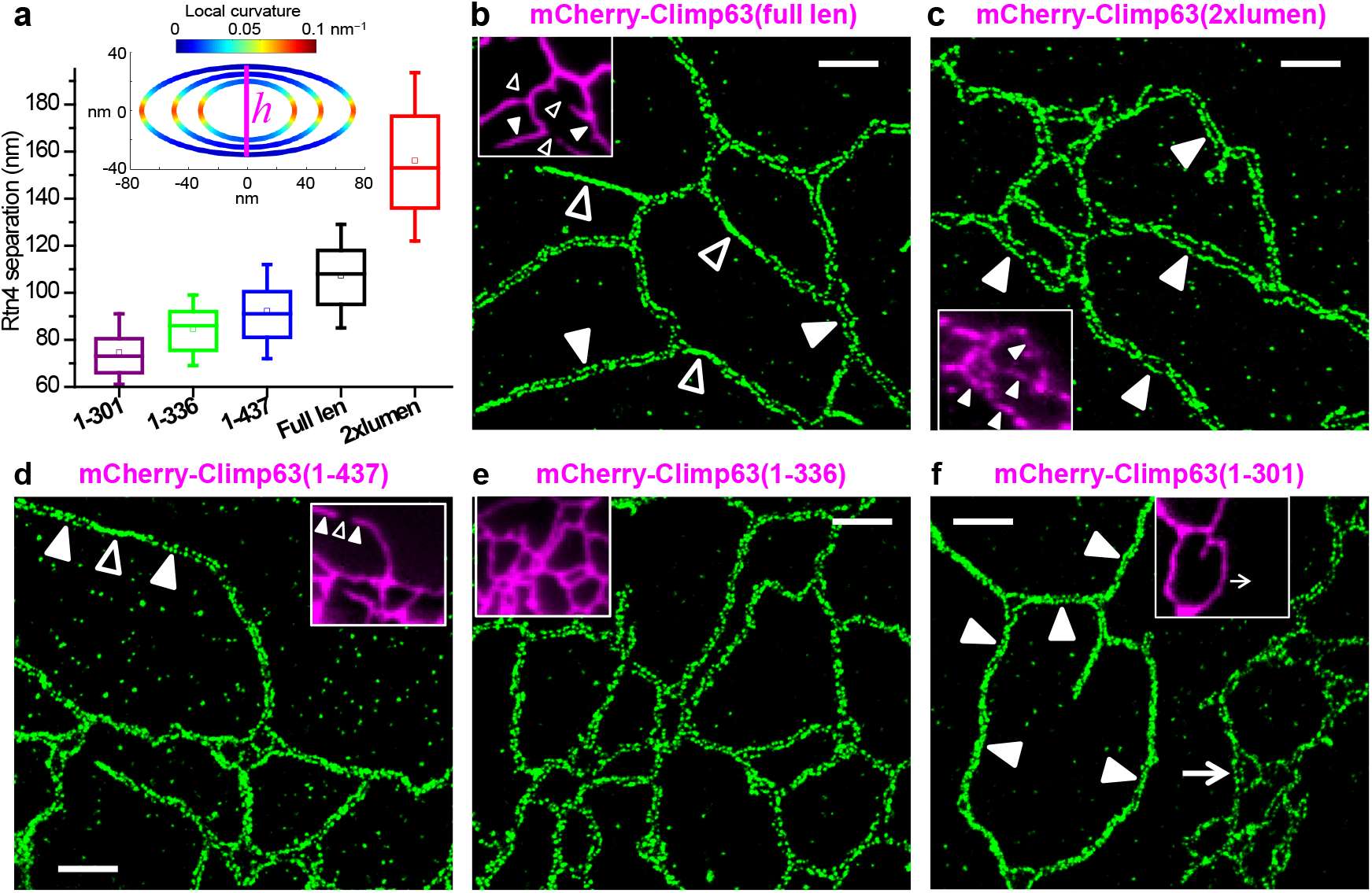
Rtn4 and Climp63 jointly define the R2 tubule width. (**a**) Distribution of center-to-center separations between opposing Rtn4 lines in R2 tubules, for COS-7 cells expressing mCherry-tagged Climp63 mutants of different intraluminal sizes. Whiskers and boxes show 10%, 25%, 50%, 75%, and 90% quantiles. For each data point, *n* = 200 local distances pooled from 5 cells. Inset: a simple model of the R2 tubule based on an elliptical cross section, in which the luminal height *h* is varied as the membrane curvature at the two edges is held at 1/*r*e = 0.08 nm-1. Color presents the local curvature. (**b-f**) Representative STORM images of the immunolabeled endogenous Rtn4 in COS-7 cells expressing mCherry-tagged full-length (1-575) (b), 2x lumen (c), 1-437 (d), 1-336 (e), and 1-301 (f) mutants of mouse Climp63. Insets: epifluorescence images of the mCherry channel. Filled and hollow arrowheads point to examples of R2 and R1 tubules, respectively. In (f), the arrow points to an untransfected cell in the same view, which facilitates a direct comparison. Scale bars: 1 µm.

### The R1 and R2 tubules differentially accommodate ER proteins

The above contrasting structures of R1 and R2 tubules may differentially regulate the distributions of ER proteins. With R1 tubules of ∼16 nm luminal diameter and R2 tubules of elliptical width and height of ∼100 and ∼50 nm, the luminal cross-sectional areas of the two tubule types differ by ∼20 folds. Further considering the protein exclusion volume, for a protein 5 nm in size, the accessible luminal volume per unit length is ∼35-fold different between the two tubule types. Indeed, two-color STORM of immunolabeled Rtn4 versus the endogenous calreticulin (Fig. 4a) and the expressed ER-residing fluorescent protein (FP) mEmerald-ER-3 (Fig. 4b) showed that both ER-luminal proteins were abundant in the R2 tubules (filled arrowheads) but absent from the R1 tubules (hollow arrowheads). Our model also predicts a ∼4-fold difference in the cross-sectional circumference, and hence the membrane surface area per unit length, between the R2 and R1 tubules. Accordingly, two-color STORM showed reduced, yet not eliminated, presence of overexpressed sec61b, a common ER-membrane marker, in the R1 tubules when compared to the R2 tubules (Fig. 4c). Interestingly, we next found that calnexin, an ER transmembrane protein with a large intraluminal domain^31^, was largely excluded from the R1 tubules (Fig. 4d). To examine the role of the intraluminal domain, we expressed in COS-7 cells two constructs with the transmembrane domain of calnexin (canxTM) linked to the 27 kDa GFP mEmerald at the intraluminal and extraluminal (cytosolic) sides, respectively. Two-color STORM with Rtn4 showed that the former was more excluded from the R1 tubules than the latter (Fig. S6). Together, our results suggest that intraluminal crowding prevents both ER-luminal proteins and ER-membrane proteins with large intraluminal domains from entering the R1 tubules. The dynamics of this regulating mechanism is further examined below with live-cell experiments.

**Fig. 4.**
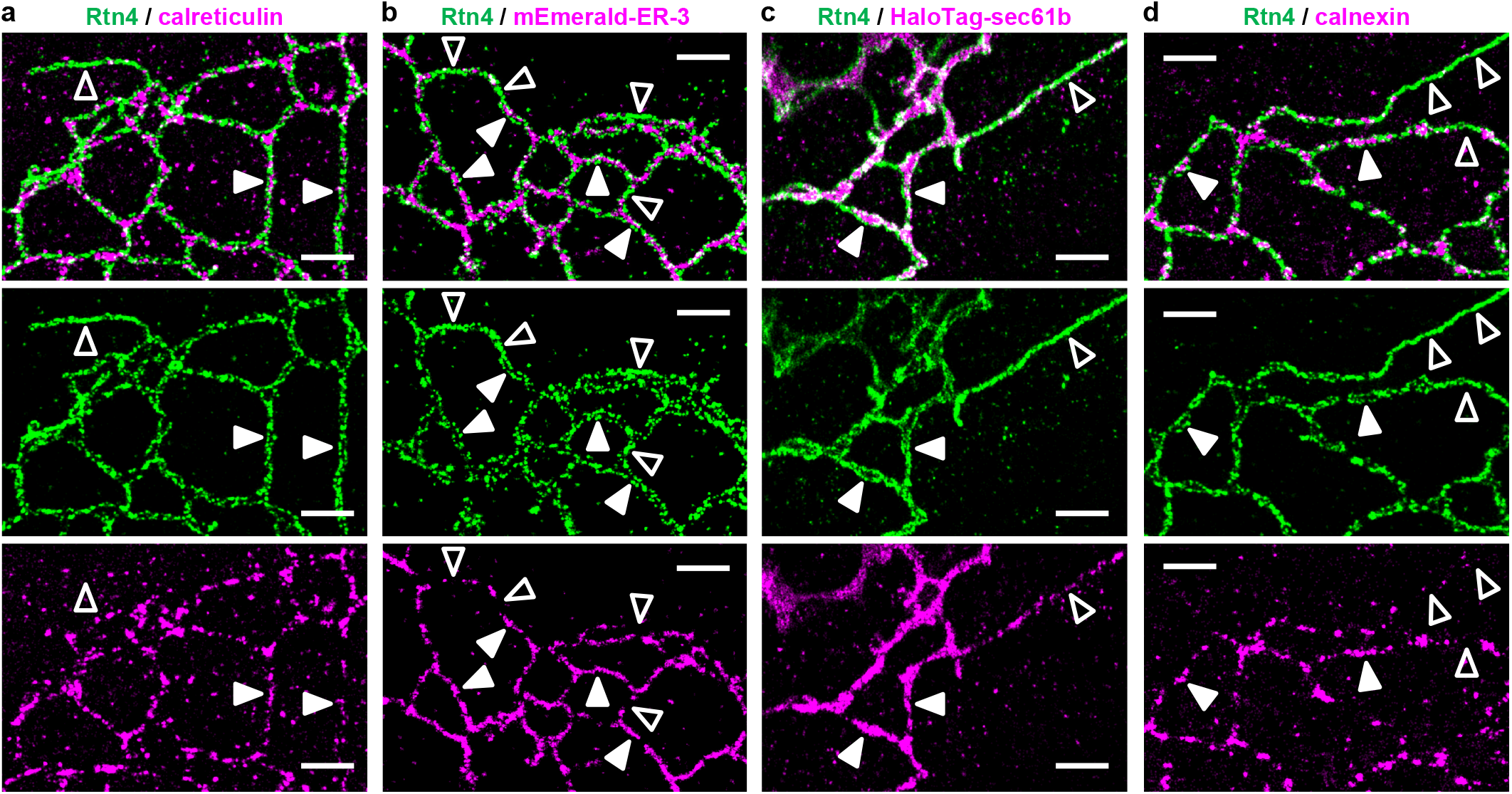
R1 and R2 tubules differentially accommodate ER proteins. (**a**) Representative two-color STORM image of the immunolabeled endogenous Rtn4 (green) and calreticulin (magenta) in an untransfected COS-7 cell. (**b**,**c**) Representative two-color STORM images of the immunolabeled endogenous Rtn4 (green) and expressed ER-luminal protein mEmerald-ER-3 [magenta in (b)] and ER-membrane protein HaloTag-sec61b [magenta in (c)] in COS-7 cells. (**d**) Representative two-color STORM image of the immunolabeled endogenous Rtn4 (green) and calnexin (magenta) in an untransfected COS-7 cell. Filled and hollow arrowheads point to examples of R2 and R1 tubules, respectively. Scale bars: 1 µm.

### R1-R2 dynamics in live cells

To probe the R1-R2 tubule dynamics in live cells, we co-expressed Rtn4b-GFP and mCherry-Climp63 in COS-7 cells. Although the single overexpression of Rtn4 and Climp63 respectively biased ER tubules toward R1 and R2 (Fig. 2bc, S3, and S4), we reasoned that the co-expression of both proteins may counterbalance the two tubule forms. Previous work has noted normal ER morphologies in cells co-overexpressing Rtn4 and Climp63^11^. We observed a mixed population of both mCherry-Climp63-positive and -negative ER-tubule segments (filled and open arrowheads in Fig. 5a). Correlating the same view with STORM of immunolabeled Rtn4 (Fig. 5b and Fig. S7) showed that the two segment forms scrupulously corresponded to R2 and R1, respectively, consistent with our model above. Thus, by tracking in the wide-field micrographs which tubule segments were Climp63-positive, we followed the R1-R2 tubule dynamics in live cells at high temporal resolutions.

**Fig. 5.**
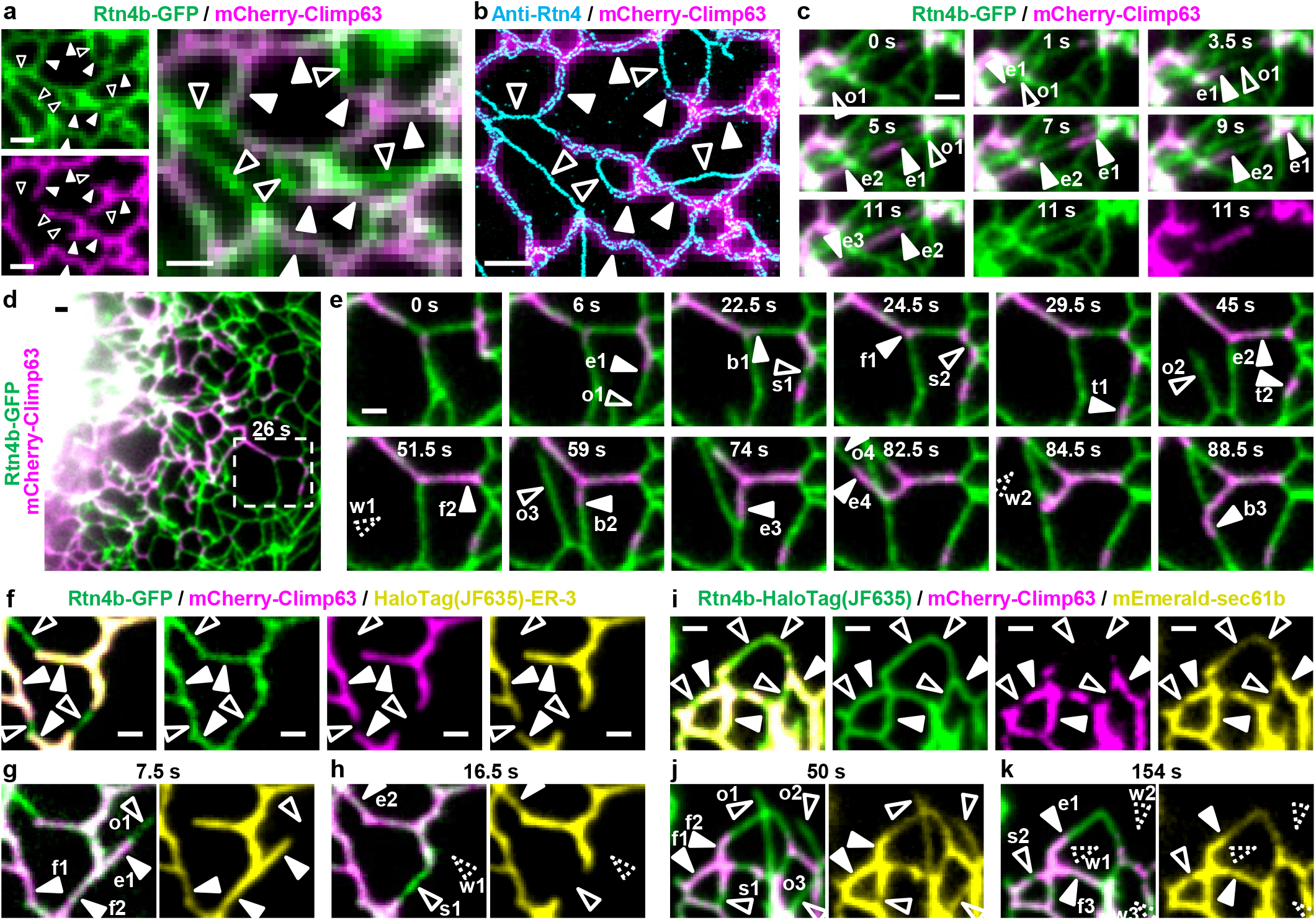
Live-cell imaging unveils fast R1-R2 remodeling and associated co-traveling of ER proteins. (**a**) Two-color fluorescence micrographs of Rtn4b-GFP (green) and mCherry-Climp63 (magenta) co-expressed in a COS-7 cell, shown as separate channels (left) and an overlay (right). (**b**) STORM of immunolabeled Rtn4 of the same view (cyan), overlayed with the mCherry-Climp63 micrograph (magenta). Filled and hollow arrowheads point to R2 and R1 examples, respectively. (**c**) Two-color micrographs of Rtn4b-GFP (green) and mCherry-Climp63 (magenta) for a small region in a living COS-7 cell at selected time points, highlighting ER-tubule outgrowth with a Climp63-free R1 tip (o1) and the ensuing extension of Climp63 into the tubule in segments (e1-e3). The two separated color channels are shown for the final image. See full series in Movie S1. (**d**) Another two-color live-cell dataset. (**e**) Selected time points for the boxed region. Arrowheads point to major structural changes vs. the preceding image. o1-o4: tubule outgrowths with Climp63-free R1 tips. e1-e4: extension of Climp63 into R1 tubules. s1-s2: the sequential splitting of a Climp63-positive R2 segment into three R2 segments connected by Climp63-negative R1 segments. f1-f2: fusion of an R2 tubule with two small R2 segments. t1-t2: translational motions of two R2 segments along tubules. b1-b3: Climp63 entering new tubule branches. w1-w2: withdrawal of newly extended tubules. See full series with separated color channels in Movies S3 and S4. (**f**) Live-cell three-color imaging of Rtn4b-GFP (green), mCherry-Climp63 (magenta), and JF635-labeled HaloTag-ER-3 (yellow) in a COS-7 cell, shown as merged and separated channels. (**g**,**h**) The same region after 7.5 s (g) and 16.5 s (h), shown as merged Rtn4b-GFP/mCherry-Climp63 (left) and HaloTag-ER-3 alone (right). See full series in Movie S5. (**i**) Live-cell three-color imaging of JF635-labeled Rtn4b-HaloTag (green), mCherry-Climp63 (magenta), and mEmerald-sec61b (yellow) in a COS-7 cell, shown as merged and separated channels. (**j**,**k**) The same region after 50 s (j) and 154 s (k), shown as merged Rtn4b-HaloTag/mCherry-Climp63 (left) and mEmerald-sec61b alone (right). See full series in Movie S6. Filled and hollow arrowheads in (f,i) point to R2 and R1 examples, respectively. Arrowheads in (g,h,j,k) point to major structural changes annotated similarly as (e). Scale bars: 1 µm.

Both the R1 and R2 tubules were highly dynamic, with extensive restructuring constantly occurring at the second/sub-second time scales (Movies S1-S4). For the fast-extending ER tubules, we found that the outgrowing tips were often in the Climp63-free R1 form (o1 in Fig. 5c; Movies S1 and S2). Climp63 then gradually extended into the tubules (e1 in Fig. 5c) to establish the R2 form, and interestingly, may do so in consecutive segments (e1-e3 in Fig. 5c; Movie S1).

A trove of intriguing R1-R2 dynamics was further observed in the tubule networks (Fig. 5de and Movies S2-S4). Besides the above-noted extension of Climp63 into R1 tubules (e1-e4 in Fig. 5e), we often observed the splitting of Climp63-positive R2 tubules into multiple R2 segments connected by Climp63-free R1 segments (s1-s2 in Fig. 5e), the fusion between R2 segments (f1-f2 in Fig. 5e), the translational motion of R2 segments (t1-t2 in Fig. 5e), the entering of R2 segments into new branches (b1-b3 in Fig. 5e), and the quick withdrawn (w1-w2 in Fig. 5e) of newly extended tubules (o2 and o4).

Three-color live-cell imaging next showed that the ER-luminal protein HaloTag-ER-3 stayed exclusively in the R2 tubules (filled arrowheads in Fig. 5f) and so co-traveled with the Climp63-positive segments throughout the R1-R2 rearrangements (Fig. 5gh and Movie S5). In contrast, the ER-membrane protein mEmerald-sec61b exhibited reduced, yet still substantial, presence in the R1 tubules (hollow arrowheads in Fig. 5i), and this distribution was also preserved as the R1-R2 tubules dynamically rearranged in the cell (Fig. 5jk and Movie S6). Comparison with STORM results on fixed cells reaffirmed the above observations (Fig. S8). Thus, consistent with our two-color STORM results above (Fig. 4), the R1 tubules excluded ER-luminal proteins but accommodated low levels of ER-membrane proteins during the fast R1-R2 remodeling.

### The R1-R2 dichotomy also applies to other ER membrane curvature proteins

Our above results with the ubiquitously abundant Rtn4 prompt the question of whether other ER-membrane curvature-promoting proteins may behave similarly. To address this issue, we expressed in COS-7 cells mEmerald-tagged Rtn3c, REEP5 (DP1), and Arl6IP1. STORM showed that when overexpressed individually, the three Rtn4-like proteins consistently covered the ER tubules in the ultrathin R1 form, with short R2 segments only sporadically spotted (Fig. 6a). These results are similar to what we observed above with Rtn4 (Fig. 2bc and Fig. S3). When co-expressed with mCherry-Climp63, all three proteins displayed R1-R2 dichotomies, so that they showed up as two parallel lines for the Climp63-positive tubule segments (filled arrowheads in Fig. 6b) but remained in the R1 form for segments devoid of Climp63 (hollow arrowheads in Fig. 6b), again mimicking the behavior of Rtn4 (Fig. 5b). These observations suggest that the R1-R2 dichotomy is a general structural arrangement for ER tubules. Different curvature-preferring membrane proteins may thus work together to both promote/stabilize the ultrathin R1 tubules and cooperate with Climp63 to stabilize the two high-curvature edges of the ribbon-like R2 tubules.

**Fig. 6.**
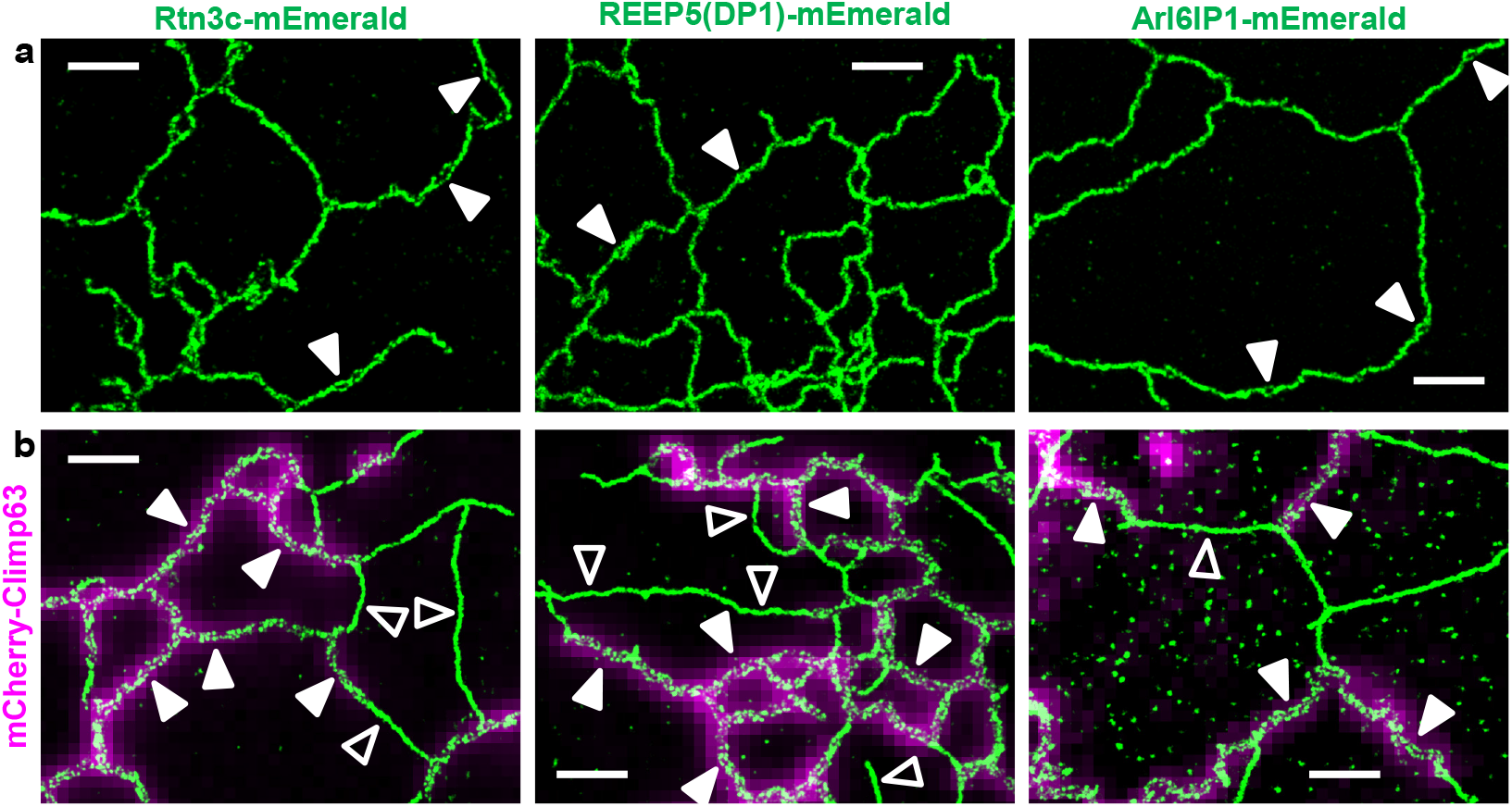
The R1-R2 dichotomy also applies to other ER membrane curvature proteins. (**a**) Representative STORM images of Rtn3c-mEmerald, REEP5(DP1)-mEmerald, and Arl6IP1-mEmerald expressed in COS-7 cells. Filled arrowhead points to small R2 segments, whereas most other tubules appear R1. (**b**) Representative STORM images of Rtn3c-mEmerald, REEP5(DP1)-mEmerald, and Arl6IP1-mEmerald (green) overlaid with epifluorescence images of mCherry-Climp63 (magenta) co-expressed in the same COS-7 cells. Filled and hollow arrowheads point to R2 and R1 examples, respectively. Scale bars: 1 µm.

## Discussion

Together, our super-resolution and live-cell microscopy results introduced an R1-R2 dichotomy for ER tubules and unveiled their dynamics and functional implications. Though our R1-R2 model is unexpected, it explains and connects several recent findings.

Live-cell STORM experiments with a membrane dye have noted that in BS-C-1 cells, the newly extended ER tubules are substantially thinner^25^. However, the origin of this phenomenon has not been elucidated. We showed that the extending “new” tubule tips were often of the thinner R1 form, yet the established “old” tubule networks were also characterized by dynamic R1-R2 rearrangements. Meanwhile, electron microscopy (EM) has visualized ∼20 nm-sized “ultrathin” ER tubules in COS cells upon Rtn4-overexpression^10^. Fluorescence microscopy notes that the overexpression of Rtn4, Rtn3c, REEP5 (DP1), Arl6IP1, and like proteins tend to “squeeze out” luminal proteins from the ER tubules^10,12,32-34^. These observations may be understood under our R1-R2 framework as that raised levels of curvature-promoting proteins drive ER tubules into the luminal-protein-excluding R1 form. Yet, our results emphasized the coexistence and dichotomy of two well-defined tubule forms in both the native and overexpressed cells. rather than continuously evolving tubule widths. Interestingly, different cell types were dominated by either of the two tubule forms, and the R1 and R2 abundances varied as we altered the intracellular levels of Climp63 and Rtn4 (and analogs). Among these results, we found that the ER tubules in the rat primary astrocytes and neurons were predominately in the R2 and R1 forms, respectively. The latter finding echoes recent EM observations that the neuronal ER tubules are curiously thin at ∼20 nm diameter^35^.

Advances in electron tomography and FIB-SEM offer more holistic pictures of the ER structure. Examination of the resultant three-dimensional models identified ribbon-like tubules^13,16,17^, including cases in which immunogold-labeled Rtn4 may have decorated the ribbon edges^16^, potentially consistent with our R2 tubule model. However, immunogold yields sparse labeling and does not readily accommodate multi-target imaging, and EM generally offers limited fields of view. STORM readily resolved Rtn4 (and analogs) throughout the cell to establish the R1-R2 dichotomy, and its multicolor capabilities further facilitated mechanistic investigations. The observed good correspondence of the Climp63-positive and -negative tubules to the R2 and R1 forms thus enabled us to employ multicolor live-cell imaging to unveil the rich second/sub-second dynamics of the two tubule forms as they differently accommodated luminal and membrane proteins.

Our results underscore how the intricate interactions between ER-shaping proteins give rise to molecularly related, yet structurally and functionally distinct ER forms. Notably, in addition to identifying a conserved R2 tubule width across different cell types, we further established a positive correlation between this width and the Climp63 intraluminal size, thus emphasizing the co-regulation of both the R1-R2 dichotomy and the final tubule geometry by Climp63 and membrane curvature-promoting proteins. How the Climp63-Rtn4 (and analog) interplays highlighted here further cooperate with other ER-shaping proteins, as well as how the R1 and R2 tubules, with their contrasting accommodation of luminal and membrane proteins, respectively participate in the diverse ER functions, present pressing questions for future experimental and theoretical efforts.

## Materials and methods

### Plasmids

FLAG-Climp63, GFP-Climp63, GFP-Rtn3c(mouse), and HA-REEP5(mouse) were gifts from Tom Rapoport^10,11^. Rtn4a-GFP (Addgene plasmid #61807) and mCherry-Climp63(mouse) (Addgene plasmid #136293) were gifts from Gia Voeltz^36^. mEmerald-sec61b (Addgene plasmid #54249), mEmerald-ER-3 (Addgene plasmid #54082), and calnexin-mEmerlad (Addgene plasmid #54021) were gifts from Michael Davidson. pcDNA3.1(+) backbone was prepared by digesting the pcDNA3.1(+) IRES GFP plasmid (Addgene plasmid #51406; a gift from Kathleen Collins) using the restriction enzymes EcoRI (ThermoFisher, FD0274) and XbaI (ThermoFisher, FD0684). Rtn4b-GFP was generated by inserting two PCR-amplified fragments from the initial part (RTN4 AA1-AA185) and the later part (RTN4 AA1005-AA1192 plus AcGFP1) of Rtn4a-GFP into the pcDNA3.1(+) backbone using Gibson Assembly (New England Biolabs, #E2611S). Rtn4b-HaloTag was generated by inserting PCR-amplified Rtn4b from Rtn4b-GFP and PCR-amplified HaloTag from pSEMS-Tom20-HaloTag (Addgene plasmid #111135; a gift from Karin Busch) into the pcDNA3.1(+) backbone using Gibson Assembly. mCherry-Climp63(2xlumen) was prepared by assembling the full-length mCherry-Climp63(mouse) with the luminal part of mouse Climp63 (AA110-AA575). The two fragments were both PCR amplified from mCherry-Climp63(mouse) and assembled by Gibson Assembly onto the pcDNA3.1(+) backbone. The codons for the first 5 amino acid of the second luminal domain was changed from ctGgaGgaGgtCcaG to ctAgaAgaAgtAcaA, which did not alter the encoded amino acids (LEEVQ). mCherry-Climp63(1-301), mCherry-Climp63(1-336), and mCherry-Climp63(1-437) were generated by inserting the PCR-amplified corresponding front parts of mCherry-Climp63(mouse) into the pcDNA3.1(+) backbone using Gibson Assembly. HaloTag-sec61b was prepared by inserting PCR-amplified HaloTag from pSEMS-Tom20-HaloTag and sec61b fragment from mEmerald-sec61b into the pcDNA3.1(+) backbone using Gibson Assembly. HaloTag-ER-3 was generated by inserting PCR-amplified calreticulin signal peptide plus the 3AA linker from mEmerald-ER-3 and HaloTag from pSEMS-Tom20-HaloTag, plus a C-terminus KDEL ER retention signal included in the primer during the PCR amplification, into the pcDNA3.1(+) backbone. mEmerald(luminal)-canxTM was prepared by inserting the PCR-amplified calreticulin signal peptide, 3AA linker, and mEmerald from mEmerald-ER-3 and the PCR-amplified calnexin transmembrane domain plus cytosolic residues (canxTM, AA482-AA592 from human calnexin) from calnexin-mEmerald into the pcDNA3.1(+) backbone using Gibson Assembly. canxTM-mEmerald(cytosolic) was generated by inserting the PCR-amplified calnexin signal peptide plus a 7AA linker from mEmerald-calnexin and the later part of calnexin-mEmerald (canxTM-14AA linker-mEmerald) into the pcDNA3.1 backbone using Gibson Assembly. Rtn3c-mEmerald, REEP5-mEmerald, and Arl6IP1-mEmerald were generated by inserting PCR-amplified Rtn3c from GFP-Rtn3c(mouse), REEP5 from HA-REEP5(mouse), and Arl6IP1 from Myc-DDK-Arl6IP1 (OriGene #RC201681), respectively, and mEmerald from mEmerald-ER-3 into the pcDNA3.1(+) backbone using Gibson Assembly. Protein-coding sequences were verified by Sanger sequencing at the UC-Berkeley DNA Sequencing Facility.

### Antibodies

Primary antibodies used: rabbit polyclonal anti-Rtn4a/b (ThermoFisher, PA1-41220), sheep polyclonal anti-Rtn4b (R&D Systems, AF6034), rabbit polyclonal anti-calreticulin (Abcam, ab2907), rabbit polyclonal anti-calnexin (ProteinTech, 10427-2-AP), rabbit monoclonal anti-sec61b (Cell Signaling Technology, 14648S), mouse monoclonal anti-α-tubulin (Sigma, T6199, DM1A), mouse monoclonal anti-Climp63 (Enzo Life Sciences, ENZ-ABS669-0100, G1/296), mouse monoclonal anti-FLAG (Sigma-Aldrich, F1804, M2), mouse monoclonal anti-GFP (ThermoFisher, A11120), rabbit polyclonal anti-GFP (ThermoFisher, A11122), and Alexa Fluor 647-conjugated rabbit polyclonal anti-GFP (ThermoFisher, A31852). For both single-color and two-color STORM, Alexa Fluor 647-conjugated secondary antibodies (Invitrogen, goat anti-rabbit IgG, A21245; donkey anti-sheep IgG, A21448) were used to label the target for imaging under 647 nm excitation. For two-color STORM, the second target was labeled by a secondary antibody (Jackson ImmunoResearch) conjugated with CF568 succinimidyl ester (Biotium, 92131) for imaging under 560 nm excitation.

### Cell culture and transfection

COS-7, BSC-1, U2OS, A7r5, NIH-3T3, and C2C12 cells (UC-Berkeley Cell Culture Facility) were maintained in Dulbecco’s Modified Eagle Media (DMEM) supplemented with 10% fetal bovine serum (FBS) at 37 °C with 5% CO_2_. Cells were cultured on #1.5 coverslips for 2∼3 days until reaching ∼70% confluency. Rat astrocytes and neurons from the E18 hippocampus (BrainBits) were plated on poly-D-lysine-coated #1.5 coverslips. Astrocytes were cultured in the NbAstro medium (BrainBits). Neurons were cultured in the NbActiv1 medium (BrainBits) for ∼14 days. Plasmid transfection was performed using Lipofectamine 3000 (Invitrogen, L3000-008) according to the manufacturer’s specifications, using ∼0.5-1 µg per well in 24-well plates (Corning, CLS3527) or Lab Teck II chambered coverglass. Transfected cells were incubated for 1-2 days before subsequent experiments. Silencer Select siRNA against Climp63 (ThermoFisher, 4392420-s21594) and scrambled Silencer Select control siRNA (GUACCAAUUCGUAAGUGUUTT; AACACUUACGAAUUGGUACTT) were transfected using Lipofectamine RNAiMAX (ThermoFisher, 13778100) according to the manufacturer’s specifications. About 50 pmol siRNA was transfected into each well in 6-well plates. The siRNA-transfected cells were cultured for 3 days before subsequent experiments.

### Immunoblotting

Suspensions of cultured cells were centrifuged at 4,200 rpm, resuspended, and washed with Dulbecco’s Phosphate-Buffered Saline (DPBS) twice before lysis. Cells were then lysed in a lysis buffer (150 mM NaCl, 50 mM Tris, 5 mM EDTA, 1% Triton-X, pH=7.5) at 4 °C for 30 min. Neurons were directly lysed in the cell culture flask with the lysis buffer and then transferred to a microcentrifuge tube. Cell lysates were centrifuged for 20 min at 12,000 rpm. The supernatant was aspirated, and Halt Protease Inhibitor (ThermoFisher, 1862209) was added. Protein concentration was determined using Pierce Rapid BCA Protein Assay Kit (ThermoFisher, A53226). 1x lithium dodecyl sulfate (LDS) loading buffer (ThermoFisher, NP0007) and 70 mM dithiothreitol were added to 20-30 µg protein sample. The sample was then incubated at 75 °C for 15 min. Electrophoresis was performed on NuPAGE 4-12% or 10% Bis-Tris Gel (Invitrogen, NP0302BOX and NP0315BOX) in 1x MOPS SDS running buffer (Invitrogen, NP0001) for ∼2 h at 90 V. Sample was then transferred to a methanol-activated polyvinylidene difluoride (PVDF) membrane (ThermoFisher, 22860) in the transfer buffer at 18 V for 50 min. The transfer buffer was prepared as 25 mM Tris base (Acros Organics, 42457-1000), 192 mM glycine (Sigma-Aldrich, G8898), and 10% v/v methanol in water. The PVDF membrane was blocked with 5% bovine serum albumin (BSA) in TBST (20 mM Tris, 150 mM NaCl, 0.1% Tween 20, pH∼7.5) for 30min at room temperature (RT). The membrane was incubated with diluted primary antibodies (1:1000, rabbit polyclonal anti-Rtn4a/b; 1:2000 mouse anti-alpha tubulin) in 5% BSA in TBST at 4 °C overnight and then washed three times with TBST buffer. The membrane was then incubated with the corresponding Alexa Fluor 647-conjugated secondary antibody or Alexa Fluor 555-conjugated anti-mouse IgG1 secondary antibody (Sigma-Aldrich, SAB4600301-50UL) at RT for 1 hour followed by 3 additional wash with TBST buffer. The membrane was then imaged with a GE Typhoon FLA 9500 Variable Mode Laser Scanner Image Analyzer.

### Immunofluorescence

Cells were fixed with 3% (v/v) paraformaldehyde (Electron Microscopy Sciences, #15714) and 0.1% (v/v) glutaraldehyde (Electron Microscopy Sciences, #16365) in DPBS at RT for 20 min, and then washed twice with a freshly prepared 0.1% (w/w) NaBH_4_ solution followed by three additional washes with DPBS. Samples were then blocked and permeabilized using a DPBS-based blocking buffer containing 0.1% (w/w) saponin (Sigma-Aldrich, S4521) and 5% (v/v) donkey serum (Jackson ImmunoResearch, 017000121) for sheep antibodies or 3% (w/w) BSA (Sigma-Aldrich, A3059) for other antibodies, for 30 min at RT. Samples were next incubated with diluted primary antibodies (1:200, sheep anti-Rtn4b; 1:100, rabbit anti-Rtn4a/b; 1:100, rabbit anti-calreticulin; 1:100, rabbit anti-calnexin; 1:100, rabbit anti-sec61b; 1:200, mouse anti-FLAG; 1:400, mouse anti-GFP; 1:200, rabbit anti-GFP) in the blocking buffer overnight at 4 °C. Samples were washed three times with washing buffer (0.1x blocking buffer diluted with DPBS) after primary labeling. Samples were then incubated with diluted dye-labeled secondary antibodies (1:400 for commercial antibodies, 1:50 for homemade antibodies) in the blocking buffer for 30 min at RT, followed by three additional washes with the washing buffer.

### STORM super-resolution microscopy

STORM imaging was performed on a homebuilt inverted microscope using a Nikon CFI Plan Apo λ 100x oil-immersion objective (NA = 1.45), as described previously^37,38^. The dye-labeled samples were mounted with a Tris-HCl-based imaging buffer containing 5% (w/v) glucose, 100 mM cysteamine (Sigma-Aldrich, 30070), 0.8 mg/mL glucose oxidase (Sigma-Aldrich, G2133), and 40 µg/mL catalase (Sigma-Aldrich, C30). Diffraction-limited wide-field images were first sequentially recorded for the different color channels using weak (∼50 mW/cm^2^) laser excitations at 647 nm (for Alexa Fluor 647), 560 nm (for mCherry and CF568), and 488 nm (for GFP) using matching bandpass filters. For STORM imaging of targets labeled by Alexa Fluor 647 and CF568, the sample was sequentially imaged using strong 647-nm and 560-nm lasers at ∼2 kW/cm^2^. The angle of incidence was slightly smaller than the critical angle of total internal reflection, thus illuminating a few micrometers into the sample. A weak (0-1 W/cm^2^) 405-nm laser was applied to assist photo-switching. The resulting stochastic photo-switching of single-molecule fluorescence was recorded using an Andor iXon Ultra 897 EM-CCD camera at 110 frames per second (fps), for a total of ∼80,000 frames per image. The raw STORM data were analyzed according to previously described methods^27,28,39^.

### Live-cell fluorescence microscopy

Cells were plated in Lab-Tek II chambered coverglass and transfected as described above. For imaging of HaloTag-labeled targets, 0.25 μM of JF635 HaloTag ligand (a gift from Luke Lavis) was added to the cell culture medium for 30 min and then rinsed off with the imaging medium. The imaging medium was DMEM containing HEPES (ThermoFisher, 21063029). Live-cell fluorescence microscopy was performed on a Nikon Eclipse Ti-E inverted fluorescence microscope using a CFI Plan Apo λ 100x oil-immersion objective (NA=1.45) with the additional 1.5x magnification on the microscope body. A multi-bandpass filter cube (Semrock Di01-R405/488/561/635 as the dichroic mirror and Chroma ZET405/488/561/640m as the emission filter) was used. Two-color imaging of GFP and mCherry was achieved by alternating the excitation laser between 488 and 560 nm in successive frames as an Andor iXon Ultra 897 EM-CCD camera recorded at 4 fps, hence an effective time resolution of 0.5 s for two-color wide-field images. Three-color imaging of GFP, mCherry, and JF635 was achieved by alternating the excitation laser between 488 nm, 560 nm, and 647 nm in successive frames at camera framerates of 4 or 3 fps, hence effective time resolutions of 0.75 s or 1 s for three-color wide-field images.

## Acknowledgements

We acknowledge support by the National Institute Of General Medical Sciences of the National Institutes of Health (DP2GM132681), the Packard Fellowships for Science and Engineering, and the Pew Charitable Trusts, to K.X. K.X. is a Chan Zuckerberg Biohub investigator.

## Supplemental information

**Fig. S1.**
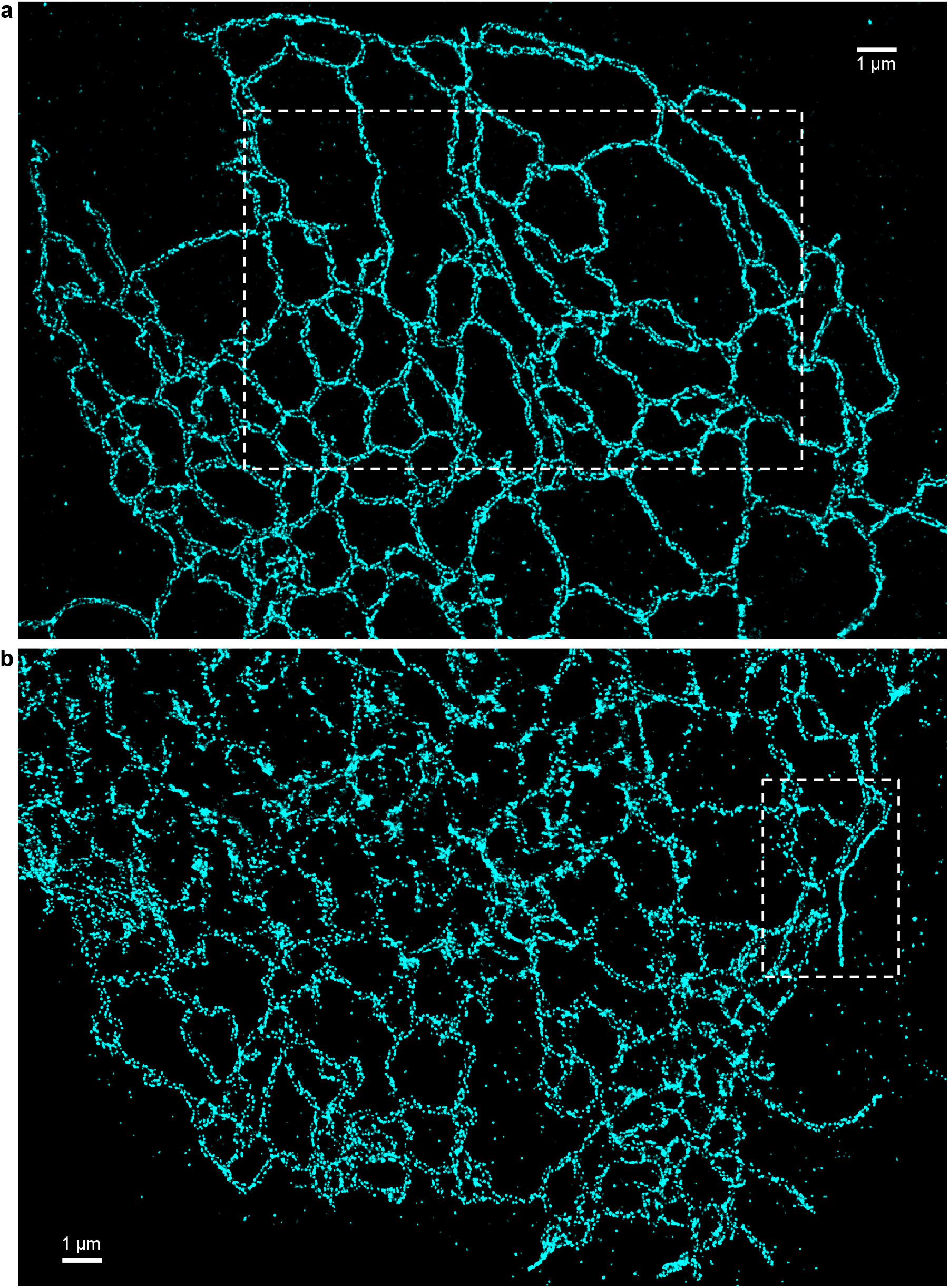
Representative STORM results of the native COS-7 cell using two anti-Rtn4 antibodies. (**a**) Cell immunolabeled with the sheep anti-Rtn4b antibody. (**b**) Cell immunolabeled with the rabbit anti-Rtn4a/b antibody. The boxed areas in (a,b) correspond to Fig. 1a and Fig. 1b, respectively.

**Fig. S2.**
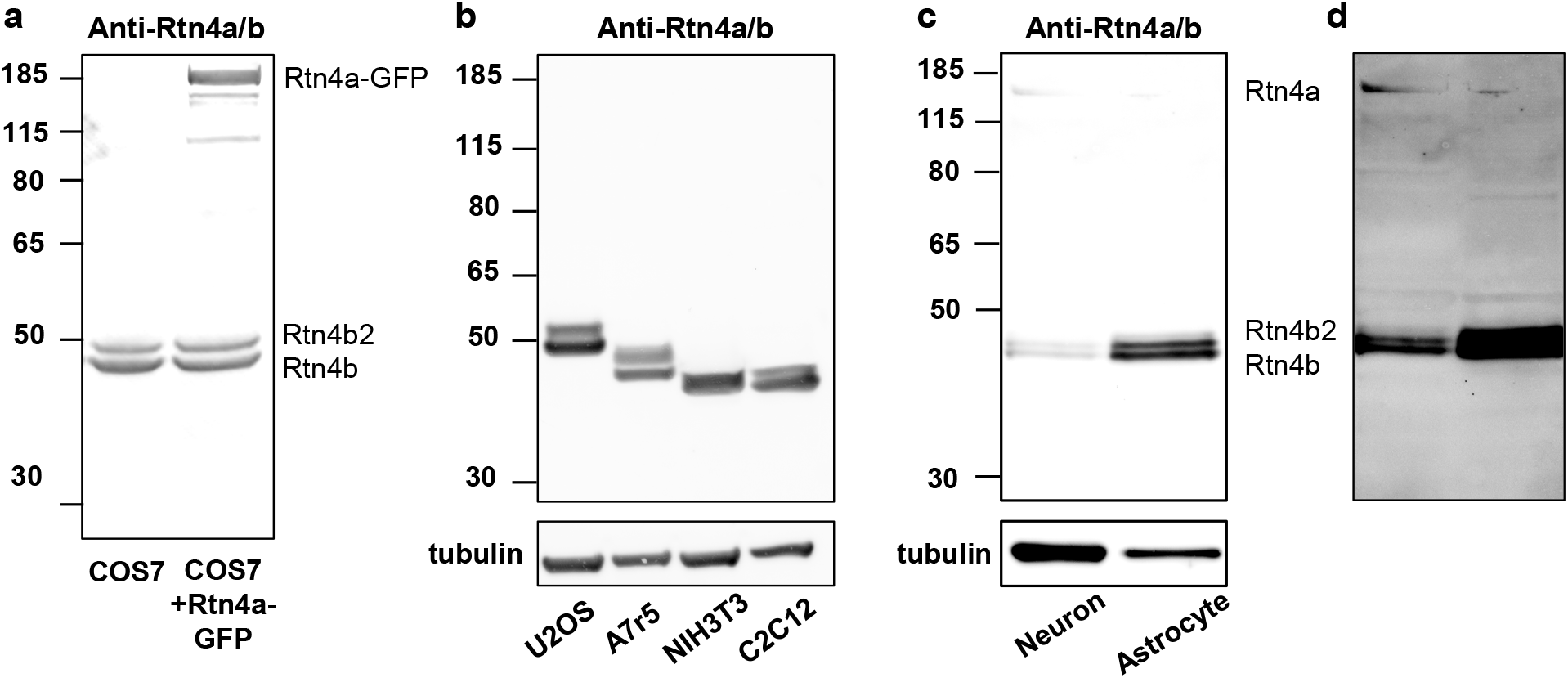
Immunoblotting results. (**a**) Immunoblotting with the rabbit anti-Rtn4a/b antibody for lysates of the native COS-7 cells (left lane) and COS-7 cells expressing Rtn4a-GFP (right lane). This showed that although the antibody also detects Rtn4a, the major forms in the native COS-7 cell are Rtn4b and Rtn4b2. (**b**) Immunoblotting with the same antibody, for lysates of untransfected U2OS, A7r5, NIH-3T3, and C2C12 cells, showing that Rtn4b and Rtn4b2 are the major forms. (**c**) Immunoblotting with the same antibody, for lysates of cultured primary neurons and astrocytes from the rat hippocampus. (**d**) Enhanced contrast for (c) to visualize the weak signal of Rtn4a, which is presumably present at the neuron plasma membrane.

**Fig. S3.**
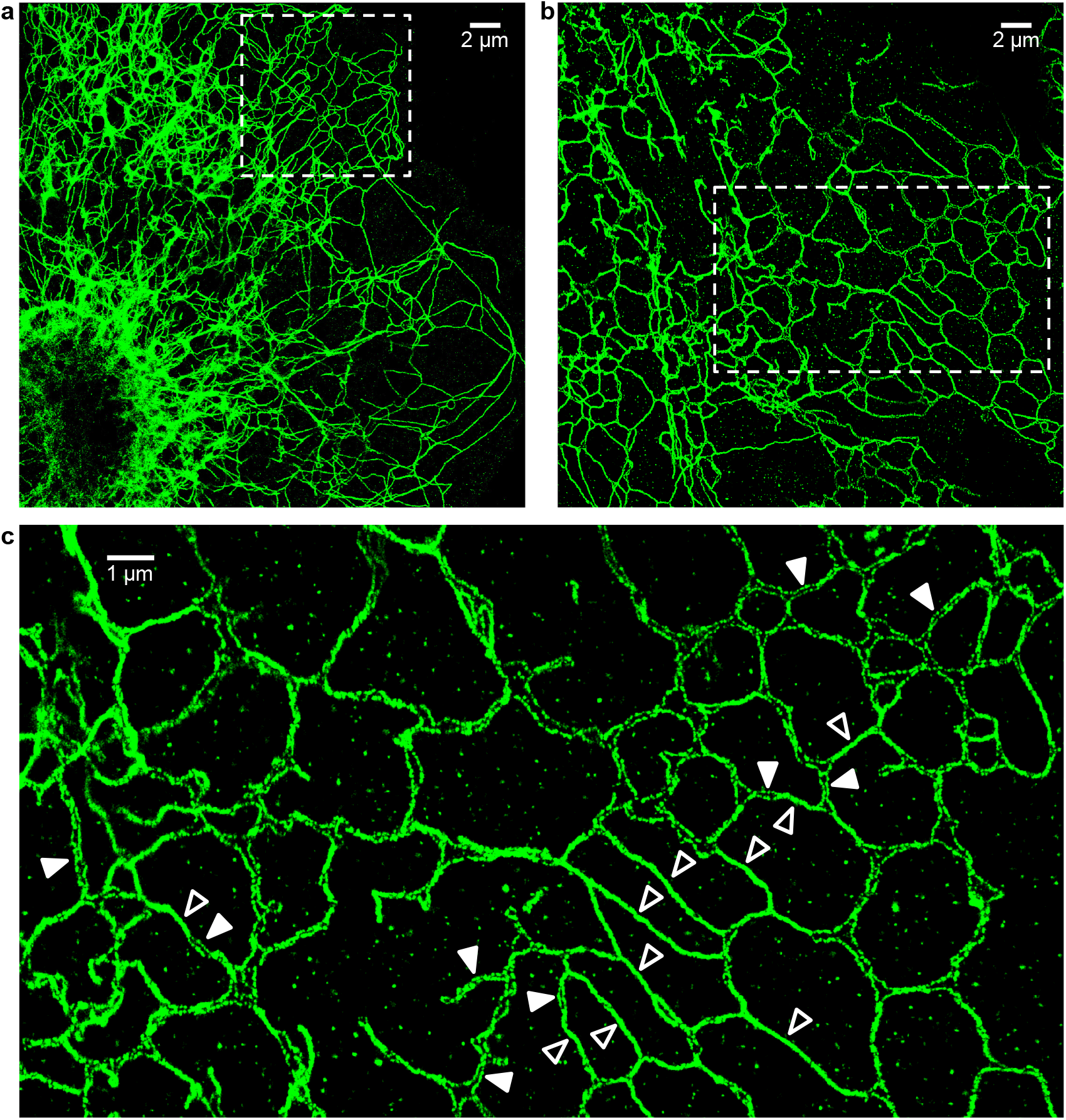
Additional results on Rtn4 overexpression. (**a**) STORM image of immunolabeled Rtn4 in a COS-7 cell overexpressing Rtn4b-GFP, showing the formation of extensive ER tubules in the R1 form. The boxed area corresponds to the magnified view in Fig. 2b. (**b**,**c**) The same conditions, but showing a cell that formed fewer ER tubules and retained more R2 tubules. (c) is a zoom-in of the box in (b). Filled and hollow arrowheads point to examples of R2 and R1 tubules, respectively.

**Fig. S4.**
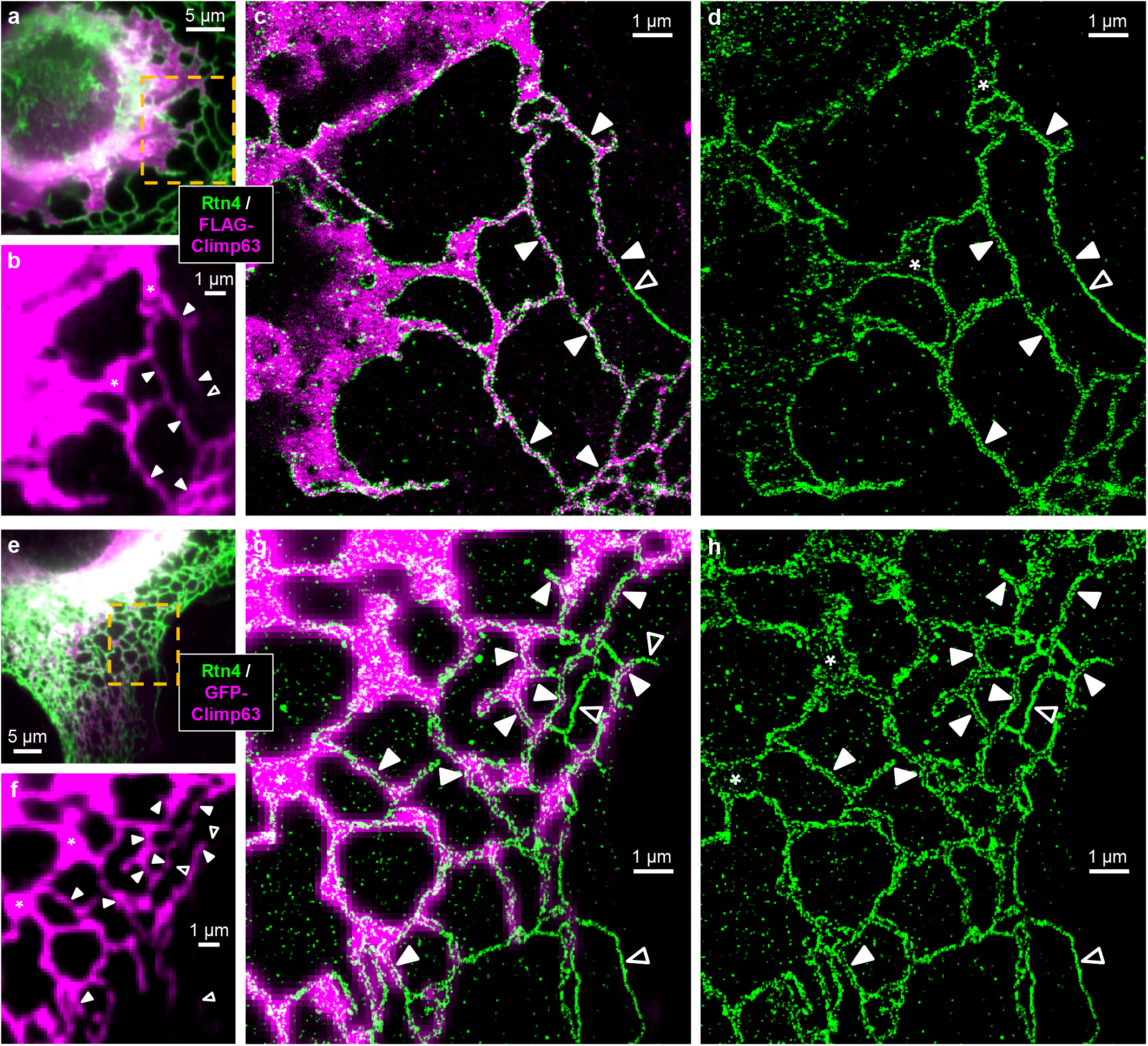
Additional results on Climp63 overexpression in COS-7 and NIH-3T3 cells. (**a-d**) A FLAG-Climp63-expressing COS-7 cell that exhibited notable expansion of ER sheets. (**a**) Two-color fluorescence image of immunolabeled Rtn4 (green) and FLAG-Climp63 (magenta). (**b**) Zoom-in of the FLAG-Climp63 channel for the boxed region in (a). (**c**) Two-color STORM image of the two channels for the same region. (**d**) The Rtn4 channel of (c). (**e-h**) An NIH-3T3 cell expressing GFP-Climp63. (**e**) Two-color fluorescence image of GFP-Climp63 (magenta) and immunolabeled Rtn4 (green). (**f**) Zoom-in of the GFP-Climp63 channel for the boxed region in (e). (**g**) The same region, overlaid with STORM image of Rtn4 (green). (**h**) The Rtn4 STORM channel of (g), showing abundant R2 tubules. Asterisks: ER sheets. Filled and hollow arrowheads point to examples of R2 and R1 tubules, respectively.

**Fig. S5.**
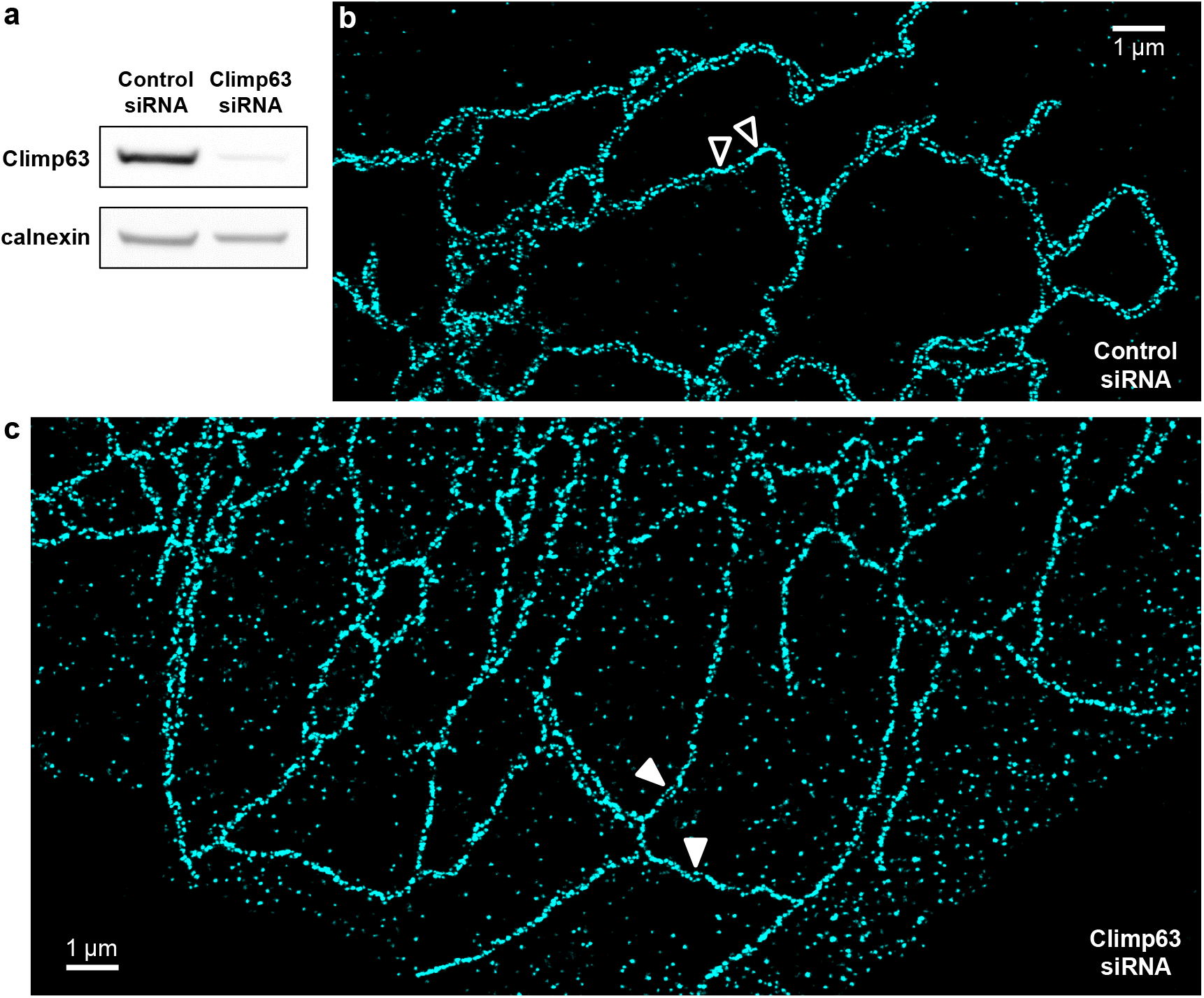
Climp63 siRNA results. (**a**) Immunoblot of Climp63 for lysates of COS-7 cells treated with control and Climp63 siRNA. Calnexin: loading control. (**b**) Representative STORM image of immunolabeled Rtn4 in a COS-7 cell treated with the control siRNA. Hollow arrowhead points to small R1 segments, whereas most other tubules in the view are R2. (**c**) Representative STORM image of immunolabeled Rtn4 in a COS-7 cell treated with the Climp63 siRNA. Filled arrowhead points to small R2 segments, whereas most other tubules in the view are R1.

**Fig. S6.**
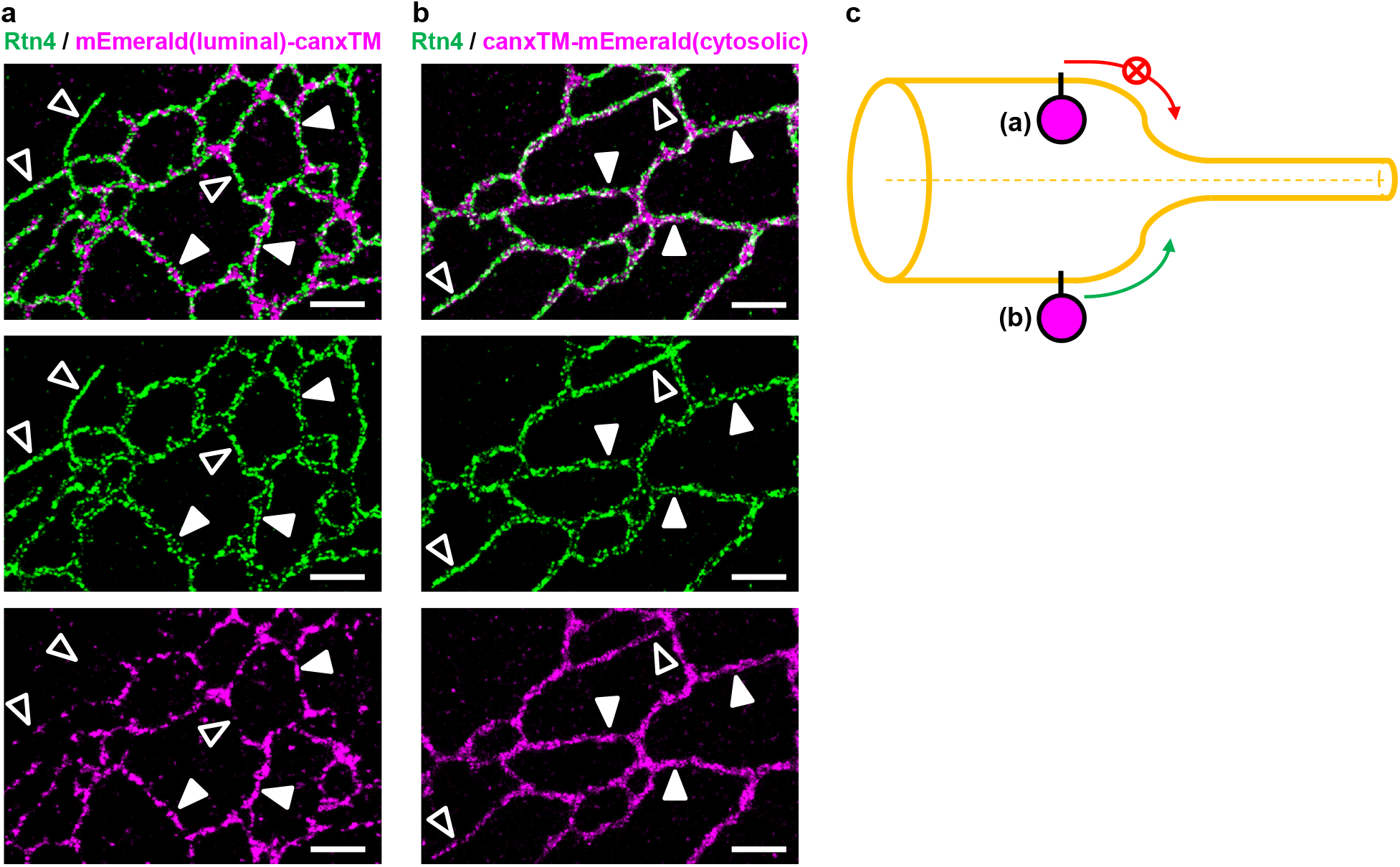
R1 and R2 tubules differentially accommodate ER transmembrane proteins with and without large intraluminal domains. (**a**,**b**) Representative two-color STORM images of the immunolabeled endogenous Rtn4 (green) and expressed calnexin transmembrane domain (canxTM) linked to mEmerald at the intraluminal (**a**) and extraluminal (cytosolic; **b**) sides, respectively, in COS-7 cells. Filled and hollow arrowheads point to examples of R2 and R1 tubules, respectively. Scale bars: 1 µm. (**c**) Model: Proteins with large intraluminal domains do not easily enter the ultrathin R1 tubule.

**Fig. S7.**
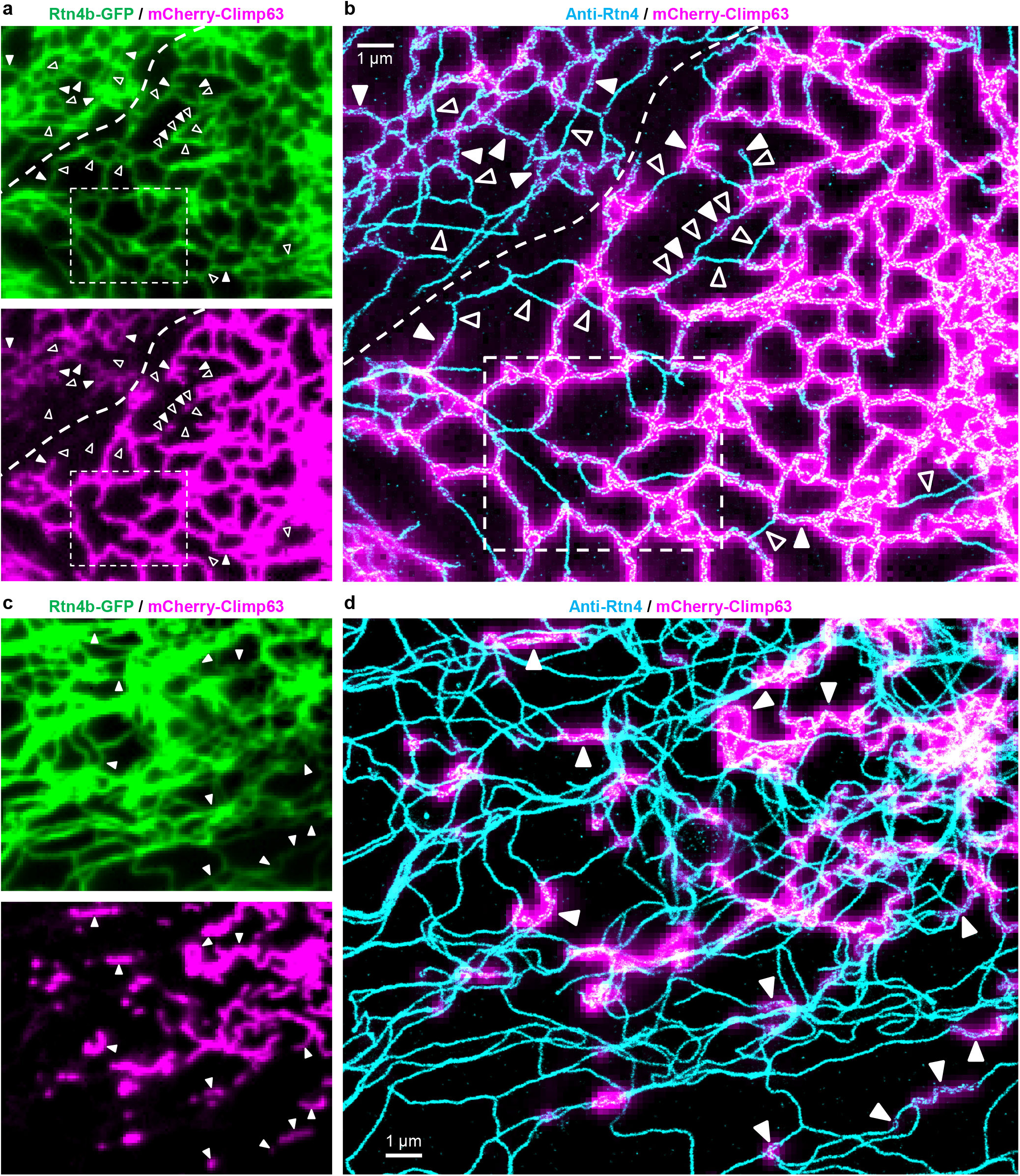
Additional results confirming that the mCherry-Climp63-positive and -negative ER-tubule segments respectively correspond to the R2 and R1 forms. (**a**) Diffraction-limited fluorescence micrographs of Rtn4b-GFP (top, green) and mCherry-Climp63 (bottom, magenta) in a co-expressing COS-7 cell. (**b**) STORM of immunolabeled Rtn4 of the same view (cyan), overlayed with the mCherry-Climp63 micrograph (magenta). The dotted curve demarcates the boundary between two cells expressing different levels of mCherry-Climp63. The boxed area corresponds to Fig. 5ab, with enhanced mCherry-Climp63 brightness to better visualize the lowly expressing cell in the upper-left corner. Both cells showed that the mCherry-Climp63-positive tubules were R2 (filled arrowheads) whereas the mCherry-Climp63-negative tubules were R1 (hollow arrowheads). (**c**,**d**) Results on another cell in which most ER tubules became R1, presumably due to the high expression level of Rtn4b-GFP. Nonetheless, the sporadic mCherry-Climp63-positive segments still correspond well to R2 (filled arrowheads).

**Fig. S8.**
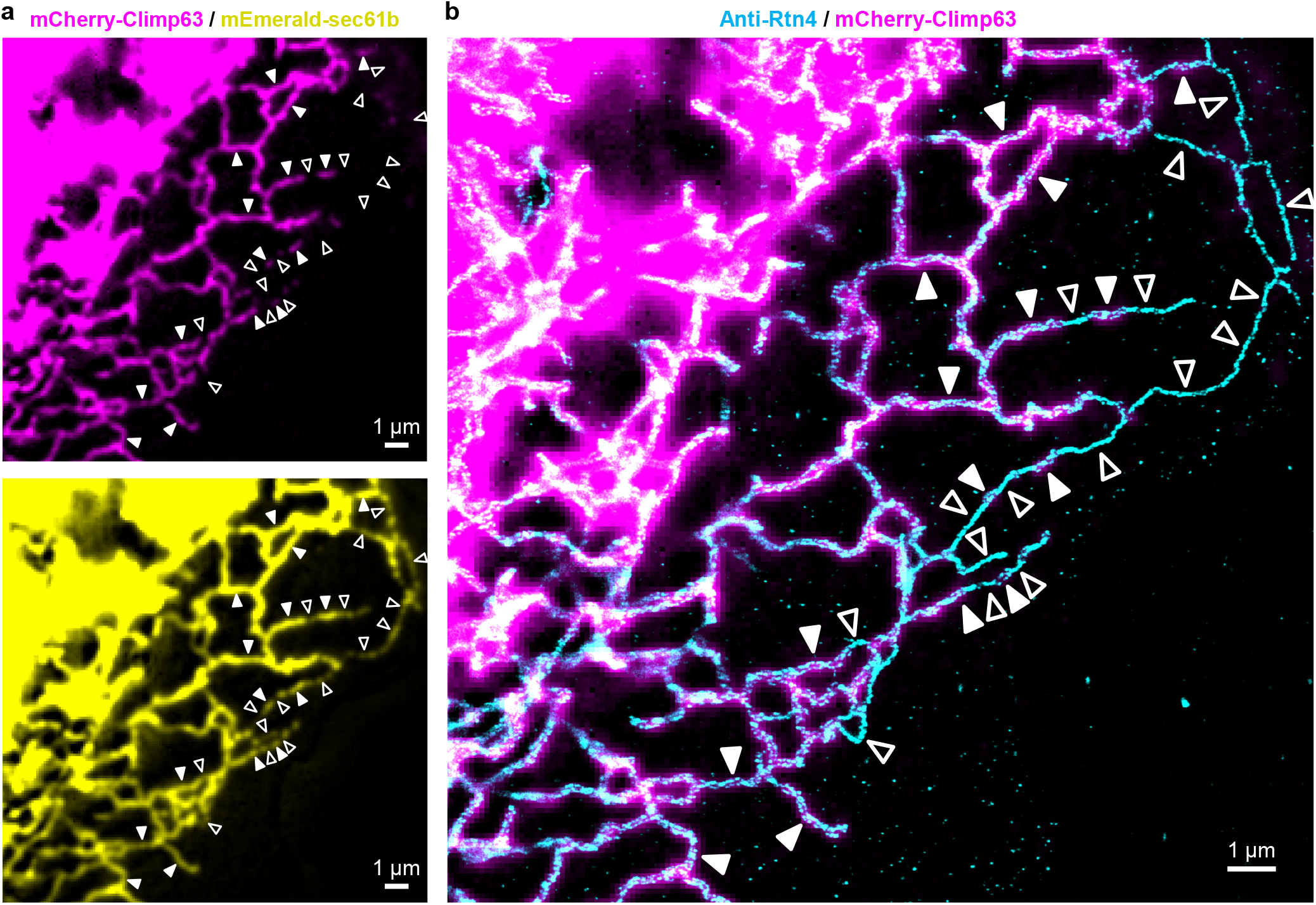
STORM of a fixed COS-7 cell co-expressing mCherry-Climp63, mEmerald-sec61b, and Rtn4b-HaloTag. (**a**) Diffraction-limited fluorescence micrographs of mCherry-Climp63 (top, magenta) and mEmerald-sec61b (bottom, yellow). (**b**) STORM of immunolabeled Rtn4 of the same view (cyan), overlayed with the mCherry-Climp63 micrograph (magenta). Filled and hollow arrowheads point to examples of R2 and R1 tubules, respectively.

## Movie captions

**Movie S1**. Consecutive time series of the two-color live-cell data shown in Fig. 5c, shown as overlaid and separate color channels for Rtn4b-GFP (green) and mCherry-Climp63 (magenta). Time 0 corresponds to the first image in Fig. 5c. Scale bar: 2 µm.

**Movie S2**. Consecutive time series from another run under the same conditions, shown as overlaid and separate color channels for Rtn4b-GFP (green) and mCherry-Climp63 (magenta). Scale bar: 2 µm.

**Movie S3**. Consecutive time series of the two-color live-cell data shown in Fig. 5e, shown as overlaid and separate color channels for Rtn4b-GFP (green) and mCherry-Climp63 (magenta). Time 0 corresponds to the first image in Fig. 5e. Scale bar: 2 µm.

**Movie S4**. Consecutive time series of the two-color live-cell data shown in Fig. 5d. Left: overlaid Rtn4b-GFP (green) and mCherry-Climp63 (magenta); Right: the mCherry-Climp63 channel alone. Time 0 corresponds to the first image in Fig. 5e. Scale bar: 2 µm.

**Movie S5**. Consecutive time series for a region containing the three-color live-cell data shown in Fig. 5f-h, shown as overlaid and separate color channels for Rtn4b-GFP (green), mCherry-Climp63 (magenta), and JF635-labeled HaloTag-ER-3 (yellow). Time 0 corresponds to Fig. 5f. Scale bar: 2 µm.

**Movie S6**. Consecutive time series for a region containing the three-color live-cell data shown in Fig. 5i-k, shown as overlaid and separate color channels for JF635-labeled Rtn4b-HaloTag (green), mCherry-Climp63 (magenta), and mEmerald-sec61b (yellow). Time 0 corresponds to Fig. 5i. Scale bar: 2 µm.

## References

1. English, A. R. & Voeltz, G. K. Endoplasmic reticulum structure and interconnections with other organelles. Cold Spring Harb. Perspect. Biol. 5, a013227 (2013).

2. Goyal, U. & Blackstone, C. Untangling the web: mechanisms underlying ER network formation. Biochim. Biophys. Acta 1833, 2492–2498 (2013).

3. Westrate, L. M., Lee, J. E., Prinz, W. A. & Voeltz, G. K. Form follows function: the importance of endoplasmic reticulum shape. Annu. Rev. Biochem. 84, 791–811 (2015).

4. Schwarz, D. S. & Blower, M. D. The endoplasmic reticulum: structure, function and response to cellular signaling. Cell Mol. Life Sci. 73, 79–94 (2016).

5. Zhang, H. & Hu, J. Shaping the endoplasmic reticulum into a social network. Trends Cell Biol. 26, 934–943 (2016).

6. Wu, H., Carvalho, P. & Voeltz, G. K. Here, there, and everywhere: The importance of ER membrane contact sites. Science 361, eaan5835 (2018).

7. Shibata, Y., Voeltz, G. K. & Rapoport, T. A. Rough sheets and smooth tubules. Cell 126, 435–439 (2006).

8. Shibata, Y., Hu, J., Kozlov, M. M. & Rapoport, T. A. Mechanisms shaping the membranes of cellular organelles. Annu. Rev. Cell Dev. Biol. 25, 329–354 (2009).

9. Voeltz, G. K., Prinz, W. A., Shibata, Y., Rist, J. M. & Rapoport, T. A. A class of membrane proteins shaping the tubular endoplasmic reticulum. Cell 124, 573–586 (2006).

10. Hu, J. J., Shibata, Y., Voss, C., Shemesh, T., Li, Z. L., Coughlin, M., Kozlov, M. M., Rapoport, T. A. & Prinz, W. A. Membrane proteins of the endoplasmic reticulum induce high-curvature tubules. Science 319, 1247–1250 (2008).

11. Shibata, Y., Shemesh, T., Prinz, W. A., Palazzo, A. F., Kozlov, M. M. & Rapoport, T. A. Mechanisms determining the morphology of the peripheral ER. Cell 143, 774–788 (2010).

12. Zurek, N., Sparks, L. & Voeltz, G. Reticulon short hairpin transmembrane domains are used to shape ER tubules. Traffic 12, 28–41 (2011).

13. Puhka, M., Joensuu, M., Vihinen, H., Belevich, I. & Jokitalo, E. Progressive sheet-to-tubule transformation is a general mechanism for endoplasmic reticulum partitioning in dividing mammalian cells. Mol. Biol. Cell 23, 2424–2432 (2012).

14. Shemesh, T., Klemm, R. W., Romano, F. B., Wang, S. Y., Vaughan, J., Zhuang, X. W., Tukachinsky, H., Kozlov, M. M. & Rapoport, T. A. A model for the generation and interconversion of ER morphologies. Proc. Natl. Acad. Sci. U. S. A. 111, E5243–E5251 (2014).

15. Wang, S., Tukachinsky, H., Romano, F. B. & Rapoport, T. A. Cooperation of the ER-shaping proteins atlastin, lunapark, and reticulons to generate a tubular membrane network. eLife 5, e18605 (2016).

16. Ramo, O., Kumar, D., Gucciardo, E., Joensuu, M., Saarekas, M., Vihinen, H., Belevich, I., Smolander, O. P., Qian, K., Auvinen, P. & Jokitalo, E. NOGO-A/RTN4A and NOGO-B/RTN4B are simultaneously expressed in epithelial, fibroblast and neuronal cells and maintain ER morphology. Sci. Rep. 6, 35969 (2016).

17. Nixon-Abell, J., Obara, C. J., Weigel, A. V., Li, D., Legant, W. R., Xu, C. S., Pasolli, H. A., Harvey, K., Hess, H. F., Betzig, E., Blackstone, C. & Lippincott-Schwartz, J. Increased spatiotemporal resolution reveals highly dynamic dense tubular matrices in the peripheral ER. Science 354, aaf3928 (2016).

18. Gao, G., Zhu, C., Liu, E. & Nabi, I. R. Reticulon and CLIMP-63 regulate nanodomain organization of peripheral ER tubules. PLoS Biol. 17, e3000355 (2019).

19. Schroeder, L. K., Barentine, A. E. S., Merta, H., Schweighofer, S., Zhang, Y., Baddeley, D., Bewersdorf, J. & Bahmanyar, S. Dynamic nanoscale morphology of the ER surveyed by STED microscopy. J. Cell Biol. 218, 83–96 (2019).

20. Shen, B., Zheng, P., Qian, N., Chen, Q., Zhou, X., Hu, J., Chen, J. & Teng, J. Calumenin-1 interacts with climp63 to cooperatively determine the luminal width and distribution of endoplasmic reticulum sheets. iScience 22, 70–80 (2019).

21. Wang, N., Clark, L. D., Gao, Y., Kozlov, M. M., Shemesh, T. & Rapoport, T. A. Mechanism of membrane-curvature generation by ER-tubule shaping proteins. Nat. Commun. 12, 568 (2021).

22. Xu, K., Zhong, G. & Zhuang, X. Actin, spectrin, and associated proteins form a periodic cytoskeletal structure in axons. Science 339, 452–456 (2013).

23. Sahl, S. J., Hell, S. W. & Jakobs, S. Fluorescence nanoscopy in cell biology. Nat. Rev. Mol. Cell Biol. 18, 685–701 (2017).

24. Sigal, Y. M., Zhou, R. & Zhuang, X. Visualizing and discovering cellular structures with super-resolution microscopy. Science 361, 880–887 (2018).

25. Shim, S. H., Xia, C., Zhong, G., Babcock, H. P., Vaughan, J. C., Huang, B., Wang, X., Xu, C., Bi, G. Q. & Zhuang, X. Super-resolution fluorescence imaging of organelles in live cells with photoswitchable membrane probes. Proc. Natl. Acad. Sci. U. S. A. 109, 13978–13983 (2012).

26. Bottanelli, F., Kromann, E. B., Allgeyer, E. S., Erdmann, R. S., Wood Baguley, S., Sirinakis, G., Schepartz, A., Baddeley, D., Toomre, D. K., Rothman, J. E. & Bewersdorf, J. Two-colour live-cell nanoscale imaging of intracellular targets. Nat. Commun. 7, 10778 (2016).

27. Rust, M. J., Bates, M. & Zhuang, X. Sub-diffraction-limit imaging by stochastic optical reconstruction microscopy (STORM). Nat. Methods 3, 793–795 (2006).

28. Bates, M., Huang, B., Dempsey, G. T. & Zhuang, X. W. Multicolor super-resolution imaging with photo-switchable fluorescent probes. Science 317, 1749–1753 (2007).

29. Beck, M., Schmidt, A., Malmstroem, J., Claassen, M., Ori, A., Szymborska, A., Herzog, F., Rinner, O., Ellenberg, J. & Aebersold, R. The quantitative proteome of a human cell line. Mol. Syst. Biol. 7, 549 (2011).

30. Schweizer, A., Rohrer, J., Slot, J. W., Geuze, H. J. & Kornfeld, S. Reassessment of the subcellular localization of p63. J. Cell Sci. 108, 2477–2485 (1995).

31. Schrag, J. D., Bergeron, J. J. M., Li, Y., Borisova, S., Hahn, M., Thomas, D. Y. & Cygler, M. The structure of calnexin, an ER chaperone involved in quality control of protein folding. Mol. Cell 8, 633–644 (2001).

32. Tolley, N., Sparkes, I. A., Hunter, P. R., Craddock, C. P., Nuttall, J., Roberts, L. M., Hawes, C., Pedrazzini, E. & Frigerio, L. Overexpression of a plant reticulon remodels the lumen of the cortical endoplasmic reticulum but does not perturb protein transport. Traffic 9, 94–102 (2008).

33. Yang, Y. S., Harel, N. Y. & Strittmatter, S. M. Reticulon-4A (Nogo-A) redistributes protein disulfide isomerase to protect mice from SOD1-dependent amyotrophic lateral sclerosis. J. Neurosci. 29, 13850–13859 (2009).

34. Yamamoto, Y., Yoshida, A., Miyazaki, N., Iwasaki, K. & Sakisaka, T. Arl6IP1 has the ability to shape the mammalian ER membrane in a reticulon-like fashion. Biochem. J. 458, 69–79 (2014).

35. Terasaki, M. Axonal endoplasmic reticulum is very narrow. J. Cell Sci. 131, jcs210450 (2018).

36. Shibata, Y., Voss, C., Rist, J. M., Hu, J., Rapoport, T. A., Prinz, W. A. & Voeltz, G. K. The reticulon and DP1/Yop1p proteins form immobile oligomers in the tubular endoplasmic reticulum. J. Biol. Chem. 283, 18892–18904 (2008).

37. Wojcik, M., Hauser, M., Li, W., Moon, S. & Xu, K. Graphene-enabled electron microscopy and correlated super-resolution microscopy of wet cells. Nat Commun 6, 7384 (2015).

38. Zhang, M., Kenny, S. J., Ge, L., Xu, K. & Schekman, R. Translocation of interleukin-1β into a vesicle intermediate in autophagy-mediated secretion. eLife 4, e11205 (2015).

39. Huang, B., Wang, W., Bates, M. & Zhuang, X. Three-dimensional super-resolution imaging by stochastic optical reconstruction microscopy. Science 319, 810–813 (2008).

